# Orientation Dependence of Current Blockade in Single Amino Acid Translocation through a Graphene Nanopore^†^

**DOI:** 10.1101/2025.03.21.644493

**Authors:** Pranjal Sur, Anurag Upadhyaya, Manoj Varma, Prabal K. Maiti

## Abstract

After successful commercialization of DNA sequencing with biological nanopores, the next frontier of the nanopore technology is protein sequencing which is far more daunting a task. Molecules passing through solid-state nanopores produce current blockades that correlate with their volume linearly in the simplest conceivable model. As thinner membranes provide better volume sensitivity, 2D materials such as Graphene, MoS_2_ membranes have been explored. Molecular dynamics studies, mostly of homogeneous polypeptide chains translocating through 2D membranes, have been reported. In this paper, we study the translocation of all the twenty single amino acids through monolayer and bilayer graphene nanopores using all-atom molecular dynamics. These studies were motivated by the fact that single amino-acids being the building blocks of peptide chains, can help us understand pore-molecule interactions during translocations at a more basic level, for instance, avoiding neighbor effects present in a chain. We show here that the correlation between the ionic current blockade and the volume of single amino acids is strongly affected by their orientation at the pore, especially when the molecule is static at pore. We explain this phenomenon by the fact that with increasing vdW volume, the amino acid in a particular orientation, has longer projection along the perpendicular direction of the pore plane. We demonstrate distinctive current and force signals for different amino-acid translocation. We observe that some of the smaller amino acids with low molecular volume produce disproportionately high current blockades in a particular orientation due to their low structural fluctuations during translocation. We investigate how dipole-moment (of the translocating amino-acids) and its alignment with electric field in the pore can be linked with our observations.

## 1 Introduction

Rapid progress in the field of single molecule sensing techniques has opened new doors for sequencing technologies such as DNA sequencing with biological nanopore^1,2^. Single-molecule protein sequencing (SMPS)^3^ is the next frontier for nanopore technology. Protein translocation and fingerprinting^4–7^ has already been reported using biological nanopores. Bio-nanopores^8^ can create a long dwell time for the translocating analyte at the pore and hence, they have been proven to be far more advantageous compared to solid-state nanopores for sequencing. Not only sequencing^9^, the nanopores can be used as a tool for understanding structure and dynamics of a complex molecule or a polymer across a confinement^10–15^.

The solid state nanopores^16–19^ are suitable for studying the structure and dynamics of proteins at a single molecule level as has been demonstrated previously^20–29^ owing to the robustness, thinness and controllability of pore properties. Hybrid pores^30–33^ that combine biological and solid-state pores are becoming potential candidates for high-fidelity detection. For solid-state nanopores, membranes with different heterostructures^34,35^ have also been experimented with. The aim of this study is to move a step towards understanding the origins of the current blockade in protein translocation via a solid-state nanopore made of graphene, starting from single amino acids and then gradually building up a larger chain to understand effects of nearest neighbors, length of the chain, conformation and dynamics of the peptide.

Previously, Molecular Dynamics studies have been performed on proteins or homogeneous and heterogeneous peptides translocating through graphene^36–39^, MoS_2_ membranes^40–43^, biological pores^44,45^ but there are no computational study involving just the single amino acids.

Here we perform MD studies of single amino acid translocation through a graphene nanopore for all twenty amino acids and compare our relative current blockades with a recent experiment^46^. Moreover, we investigate whether there exists any dependence of the orientation in the current blockade even for a molecule as small as an amino acid. We observed successful translocation events of single amino acids, indicated by a prominent dip of the processed signals near the pore. The change in current produced by each molecule is proportional to the excluded volume of that molecule^47^. The simplest conceivable model is the following.

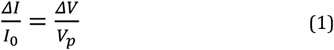

where I_0_ is the baseline current, ΔI is the deviation of the current from baseline in presence of the passing molecule, 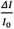 is the relative current blockade, ΔV is the excluded volume of the passing molecule and V_*p*_ is the volume of the pore. Previous studies^38,39^ have indeed shown that the amount of current reduction is correlated with the volume of the amino acids in a linear way for a homopeptide chain. We show that while this holds for even single amino acid molecule, it is dependent on the orientation of the amino acids at the pore. We performed two types of amino acid translocation studies in presence of a constant bias voltage. First, we kept each amino acid fixed or static at different orientation at the pore for the whole simulation time, and second, we pulled that amino acid (constant velocity pulling using SMD) in those same initial orientations through the pore to mimic the translocation process. We also compare data for both the monolayer and bilayer graphene pore systems. All the simulation protocols have been described in Methods section. In Fig. 1, we show our simulation setup describing the three orientations we used for amino acids and the pulling schematic. The twenty amino acids have been categorized in mainly four classes:

**Fig. 1.**
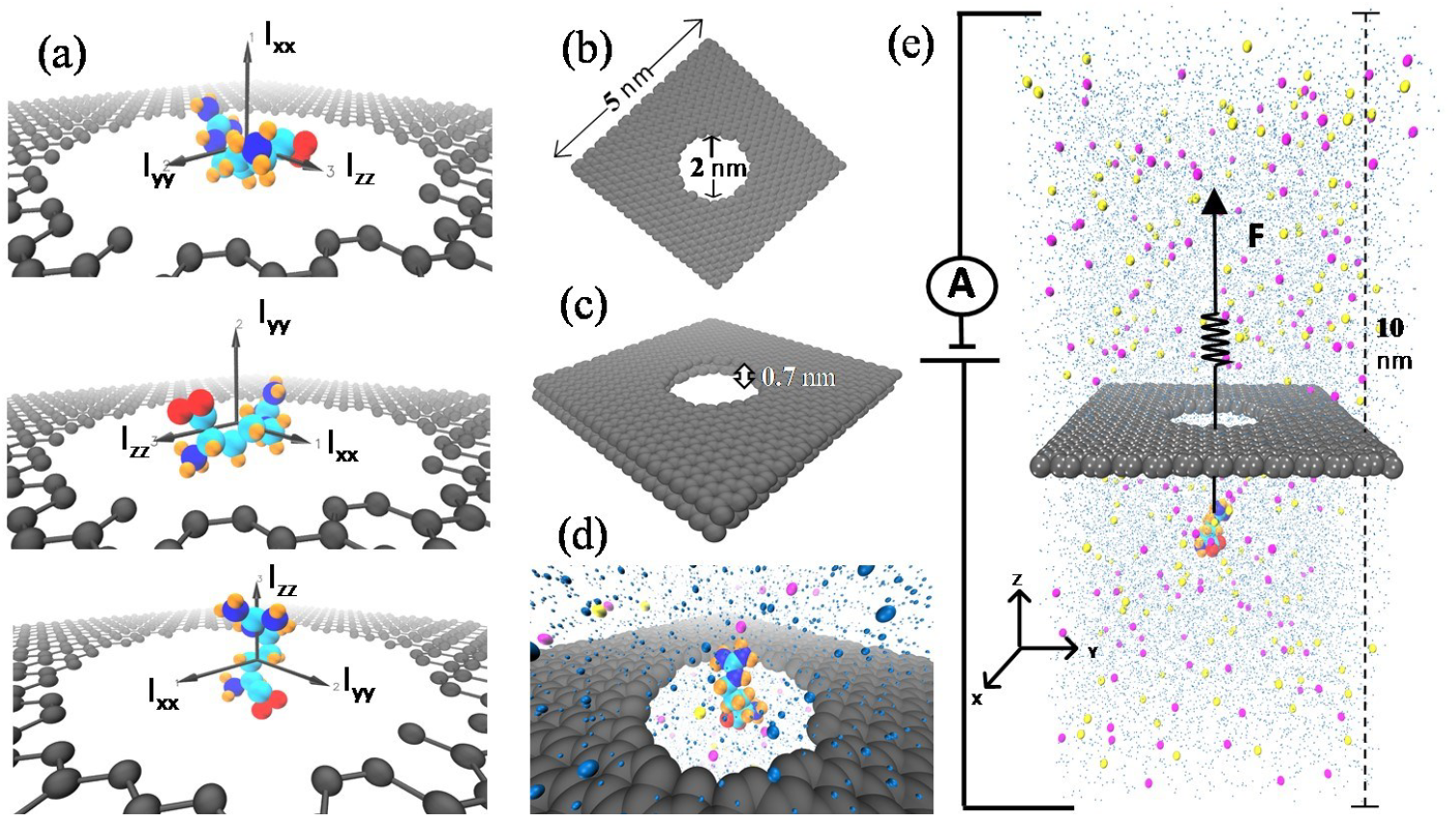
(a) Three different orientations of amino acids at the pore mouth. In each orientation, one principal moment of inertia is directed in +ve Z axis and the other two are in XY plane, (b) Pore geometry in the monolayer graphene sheet with circular pore, (c) Pore geometry in the bilayer graphene sheet with circular pore, (d) Amino acid kept static at the center of the pore, (e) Amino acid being pulled along +ve Z direction in the constant velocity ensemble.

i)**Charged and Hydrophilic** : Aspertic Acid (ASP/D), Glutamic Acid (GLU/D), Lysine (LYS/K), Arginine (ARG/R) ii)**Neutral and Hydrophilic** : Serine (SER/S), Threonine (THR/T), Asparagine (ASN/N), Glutamine (GLN/Q), Histidine (HIS/H) iii)**Neutral, Hydrophobic and Aromatic** : Phenylalanine(PHE/F),Tyrosine(TYR/Y),Tryptophan(TRP/W) iv)**Neutral**,**Hydrophobic and Non-aromatic** : Glycine (GLY/G), Alanine (ALA/A), Cystine(CYS/C), Proline(PRO/P), Leucine(LEU/L), Isoleucine (ILE/I), Methonine (MET/M), Valine(VAL/V)

Principal axes (*I*_*xx*_, *I*_*yy*_, *I*_*zz*_) for each amino acid were calculated and each principal axis was then aligned in +ve Z direction to get three different orientation (Fig1 **a**) of each amino acid. For example, if *I*_*xx*_ is aligned along Z, then that orientation is stated here as zx and has been used as a suffix (e.g., ARG_*zx*_ means Arginine in zx orientation). Similarly, *I*_*yy*_ aligned in Z is the zy orientation and *I*_*zz*_ aligned in Z is the zz orientation.

## 2 Methods

Using the nanotube builder plugin from VMD^48^, monolayer sheet (5.03 nm × 4.96 nm) and bilayer (AB stacked) sheet (armchair) of graphene was generated. A circular pore of 1 nm radius was created by removing atoms satisfying *x*^2^ +*y*^2^ < 100(Å) where *x* and *y* are coordinates of graphene atoms. Intra and inter-molecular interactions involving amino acids and graphene were described using CHARMM36m^49^ Force Field. For graphene, we use the CA (aromatic carbon) atom type in CHARMM36m as was used in previous studies^37,39^. The water model used here is the CHARMM modified TIP3P^50^. For ions, we use the parameters developed by Beglov and Roux^51^. The topology and the initial structure of the system was generated at first. Principal axes for each amino acid were calculated in VMD using the Orient package. Each principal axis was then aligned in +ve Z direction to get three different orientation of each amino acid. Then, the sheet and the molecule (for open-pore characteristics, without the molecule) was solvated with water molecules in a box of size 5.3 nm × 5.2 nm × 10 nm where the sheet is in the central plane of the box. Followed by solvation, ions were added to the system, by removing some water molecules randomly and replacing them with K^+^ and Cl^−^ ions to generate 1M KCl solution in which the sheet with the pore and molecule is immersed. For charged amino acids, the charge neutrality of the whole system was maintained by adding necessary counterions. All atom molecular dynamics simulation was performed using GROMACS 22.1^52^ and the visualization tool used was VMD^48,53^. Energy minimization of the whole system was done with the steepest descent algorithm with 0.01 nm step size till the minimization process reached the machine precision. During energy minimization, there was no restraint on amino acids. Consequently, there was a NVT equilibration followed by a NPT equilibration. For equilibrations, amino acids were restrained (C.O.M. of heavy atoms, i.e. every atom except hydrogen) in X, Y, Z with 1000 kJ mol^−^1nm^−^2 in each direction. The production run was performed in NVT. For static amino-acid studies, restraints on the amino acids in the production run were same as equilibration. But in pulling studies, restraint in Z axis was removed and restraints in X and Y were kept to resist random movement of small amino acids in order to achieve the translocation data in a reasonable runtime. In the production run, constant electric field was applied across the box along Z axis (+0.1 V/nm for amino acid studies). The voltage applied was calculated considering |*V*| = E × L_z_ where L_z_ is the box length in z dimension. A total of 1V bias was applied for all the data shown here. The sheet was restrained with a high value of 50000 kJ mol^−1^nm^−2^ throughout the whole process to resist bending of the planer sheet. For static amino-acid studies, amino acids were kept fixed at center where in pulling studies, they were pulled along Z axis from 2 nm below the pore in the direction (+ve Z) of applied electric field. For constant velocity pull, pulling velocity was 1.25 Å/ns and the spring constant was 5000 kJ mol^−1^nm^−2^. Trajectory was dumped after each 2 ps. Integration time step was 2fs and a Leapfrog integrator was used. The total inter and intra-molecular interaction energy can be described by

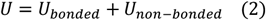

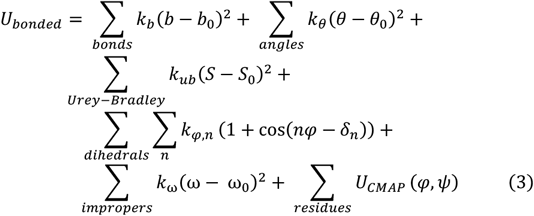

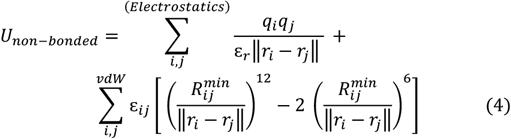

Here *b*, θ, *S*, φ, ω represents the bond lengths, valence angles, Urey-Bradley 1,3-distances, dihedral angles and improper torsion angles, respectively while *b*_0_, θ_0_, *S*_0_, ω_0_ are corresponding equilibrium values and *k*_*b*_, *k*_θ_, *k*_*ub*_, *k*_*φ*_ are corresponding force constants. In the dihedral term, *k*_*φ,n*_ is the amplitude, *n* being the multiplicity and *δ*_*n*_ is the associated phase. U_*CMAP*_ is for backbone torsion corrections. The partial atomic charges of the atoms i, j are *q*_*i*_, *q*_*j*_ while the dielectric constant is *ε*_*r*_. For particle i, ε_*i*_ is the depth of the Lennard-Jones potential well, and 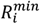 is the equilibrium radius of the *i*^*th*^ particle. The van der waals interaction between atoms i, j was modelled as per the Lennard-Jones potential where ε_*ij*_ is the effective depth of the potential well, 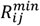 is the effective equilibrium distance. The effective parameters were calculated using the Lorentz-Berthelodt combination rules, in which ε_*ij*_ is the geometric mean of ε_*i*_ and ε_*j*_ while 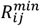 is the arithmetic mean of 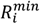 and 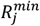. ∥*r*_*i*_ −*r*_*j*_∥ represents the Euclidean distance between atoms i, j.

SETTLE^54^ was used for rigid water model and hydrogen bond constraint was implemented with LINCS^55^ algorithm. In order to calculate vdW interactions, we have used Verlet^56^ neighbor list scheme. We used a switching function between an inner cutoff 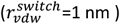 and outer cutoff radius (r_*vdw*_ =1.2 nm) to calculate the LJ potential and the forces derived from the LJ potential. This ensures a smooth transition to zero avoiding artifacts due to force discontinuity. The cutoff radius was set to be 1.2 nm for calculating the short-range part of the electrostatic interactions in real space. The long-range part of the electrostatic interactions was calculated using the Particle Mesh Ewald^57^ method. To maintain constant temperature of 300K we used the Bussi-Donadio-Parrinello stochastic velocity rescaling^58^ thermostat. The time constant for the thermostat was set to be 0.1 ps. Pressure coupling (NPT equilibration) was implemented with Berendsen^59^ barostat. The reference pressure and time constant for coupling was set to be 1 bar and 2 ps respectively. Periodic Boundary Condition was applied along all the directions. For each frame, current was calculated using,

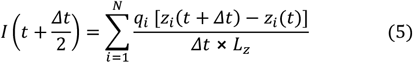

with periodic boundary corrections as had been done in previous works^21,31,39^. Here, *z*_*i*_ stands for z component of position of *i*_*th*_ ion, *q*_*i*_ is the charge of that ion, *t* is time of the frame in calculation, Δ*t* is the time gap between each frame, *L*_*z*_ is the length of the box in Z direction and *N* is the total number of ion in the frame. For processing the raw signal, block averaging was done with a 100 ps window followed by a moving average of 1.5 ns window. Subsequently, smoothing was done using Sav-Gol^60^ filter with a filter window of 21 and polynomial of order 3. After generating processed data, an average curve for both current and force had been produced by taking average of five runs in each point of time and that curve has been used for the results. For every average window, the average value is assigned to the starting time of the window, instead of assigning the value to every time stamp of that window to avoid stepwise curve. Hence, if the original trace had 20000 points, the processed trace has approximately 400 data points. For static amino-acid studies, the current for each amino acid was the simple time average of the average curve and baseline current is considered as the open-pore current. For pulling studies, average of the region |*Z*_*COM*_| = 1nm has been taken for the values of force (modulus of force), current, R_g_ and its projections in the translocation region where *Z*_*COM*_ is the z co-ordinate of center of mass of the translocating amino acid. The baseline values (when the molecule is away from the pore) in this case are the average of the region |*Z*_*C*.*O*.*M*._|> 1nm. Radius of Gyration and its projections have been calculated using the module gyrate from GROMACS. Angle fluctuations have been calculated using the GROMACS module gangle. For a parametric estimation, we fitted the angle-histogram with gaussian mixture model (GMM) to roughly represent the multi-modal cases. Here we use mixture of three gaussians,

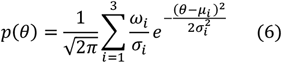

where p(θ) is the probability of angle (θ) occupation, µ_i_, σ_i_ ω_i_ are parameters assigned to mean, standard deviation and mixture weightage of gaussian components. The dipole moment, its components and its alignment with the Z axis have been calculated using the gromacs module dipoles.

## 3. Results and Discussion

### 3.1 Amino Acids Static at the Pore Center

In Fig. 2, we see that although zx and zy orientation show a linear trend between relative current blockades and volume but in zz orientation that trend is significantly reduced. As evident from the Pearson correlation coefficient shown in Fig. 2, zx orientation has the highest correlation and zz has the least. This clearly indicates that the origin of the current blockade is not solely dependent on the volume of the translocating molecule. For molecules even as small as a single amino acid, surprisingly we see orientation playing an important role. The correlation co-efficients are also different for monolayer and bilayer graphene. We see that the amino acid SER has the lowest average blockade current in five among the six panels in Fig. 2 although GLY is the lowest in volume. Even though TRP has the highest volume, it does not always produce the highest current blockade. Current blockade values of the isomers LEU and ILE are distinguishable despite having the same vdW volume, except the zz orientation (Fig. 2**c, d**). We see the existence of an overall positive correlation between blockade current magnitude and the volume of the amino acid molecule, although, there are significant deviations caused by other factors such as orientation.

**Fig. 2.**
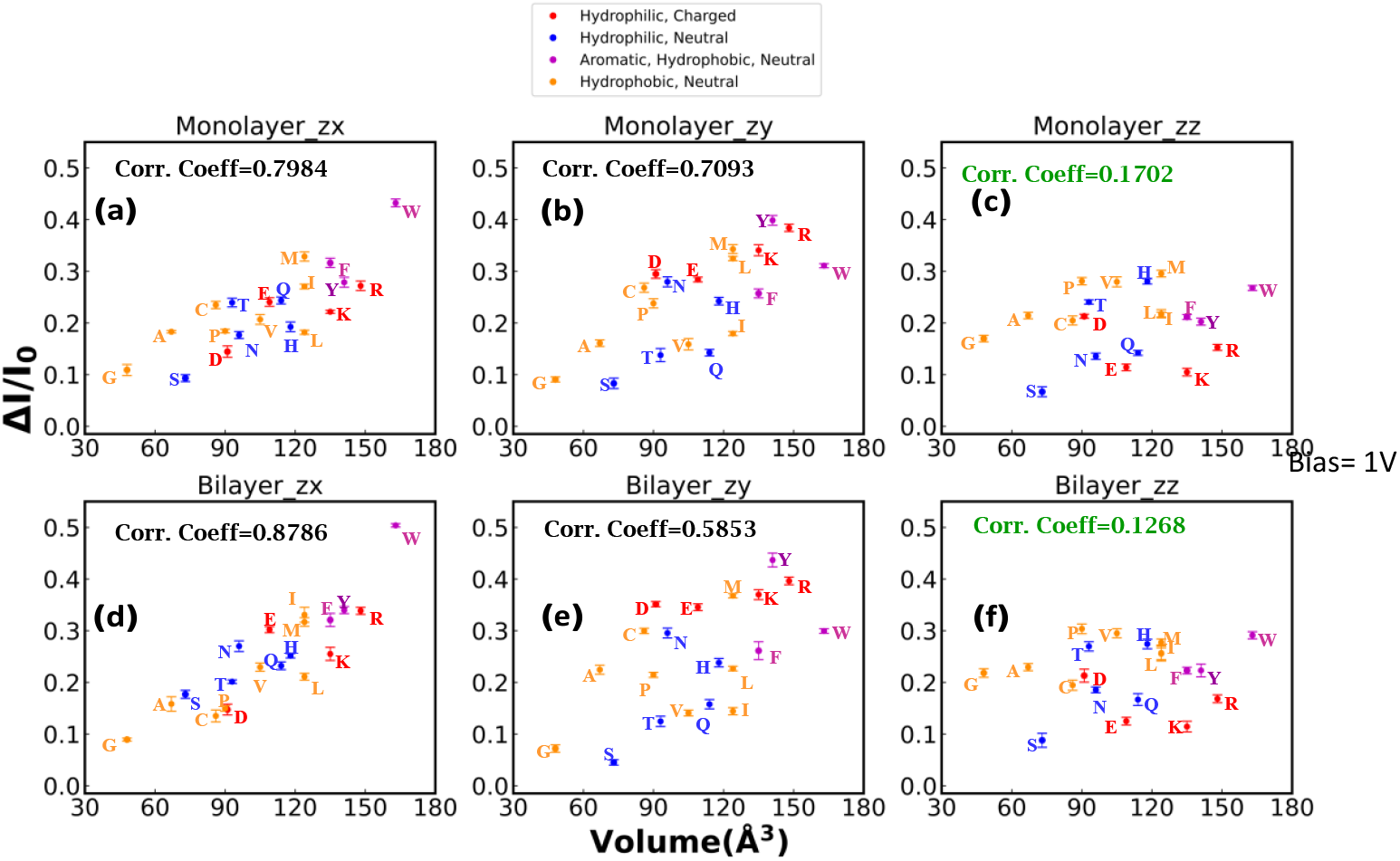
Correlation of vdW volumes and relative current blockade produced by amino acids static at pore center. Top and bottom row represents monolayer (a, b, c) and bilayer graphene pore geometry (d, e, f), respectively. The first, second and third column represents the zx (a, d), zy (b, e), zz (c, f) orientation respectively. A reduction of correlation can be seen in the third column (c, f), i.e. zz orientation. Error bars were produced using five replicas of the production run.

### 3.2 Amino Acids Translocating Through the Pore

Each of the 20 amino acids were initially kept at 2nm below the pore mouth in three different orientations and using SMD simulations they were pulled in +ve Z direction with a constant velocity. Fig. 3 describes how relative current blockades for amino acids change with their volume. From the Pearson correlation coefficient, it can be realized that correlation between volume and relative current blockade is dependent on how the amino acid is oriented at the pore and dependent on layer numbers of the membrane. The correlation is minimum for zz orientation. However, the correlation decrement is not as drastic as for the static case. In other words, while pulling, the orientation effect tends to diminish. The lowest current blockade is produced by the one least in volume, GLY except at zz orientation (bilayer) where SER is the lowest by a very small margin. In zx and zy configuration, the highest blockade is produced by a member of the high-volume aromatic, hydrophobic, neutral class of the amino acids (not strictly following order of the volume in that group) which was also the case for static amino acids. In Fig. 4, a distinct change in current and force signals can be observed as the amino acid translocate through the pore. The current blockade and the force profile for the charged amino acids in orientation zx is represented in Fig. 4**a**. Similar data for rest of the amino acids can be found in SI (Section 9, **Fig. S15-34**). Current through bi-layer pore is distinctively lower than monolayer which is consistent with the fact that bi-layer pore produces a lower open-pore current than mono-layer pore (**Fig. S1**). The maximum dip in the current does not exactly happen when COM of the amino acid coincides with center of geometry of the pore. This suggests that the amino acid, while translocating, is being stalled slightly above or below the pore plane depending on the structure and charge of that amino acid. For negatively charged ASP and GLU, the force curve is positive or rather repulsive in nature and for positively charged LYS and ARG, the force is negative or attractive. This is expected since the electric field is along +ve Z axis while translocation is in the same direction, the field helps the positively charged molecules to overcome the confinement barrier and resists the negatively charged molecules to move along positive Z. In Fig. 4**b**, translocation of two neutral amino acids have been demonstrated. For charged amino acids, the force curve is either majorly attractive or repulsive where for neutral amino acids the force curve contains both a repulsive (positive force) and attractive (negative force) region. We demonstrate in panel **c** and **d** of Fig. 4, distinctive translocation profile of current and force respectively, for the hydrophobic, neutral, non-aromatic class of amino acids in zz orientation. The probability density plot of the currents shown in Fig. 4**c** can be found in SI (section 16, **Fig. S45**). This distinctive nature is present amongst the members of other classes of amino acids (SI section11, **Fig. S36-39**) too.

**Fig. 3.**
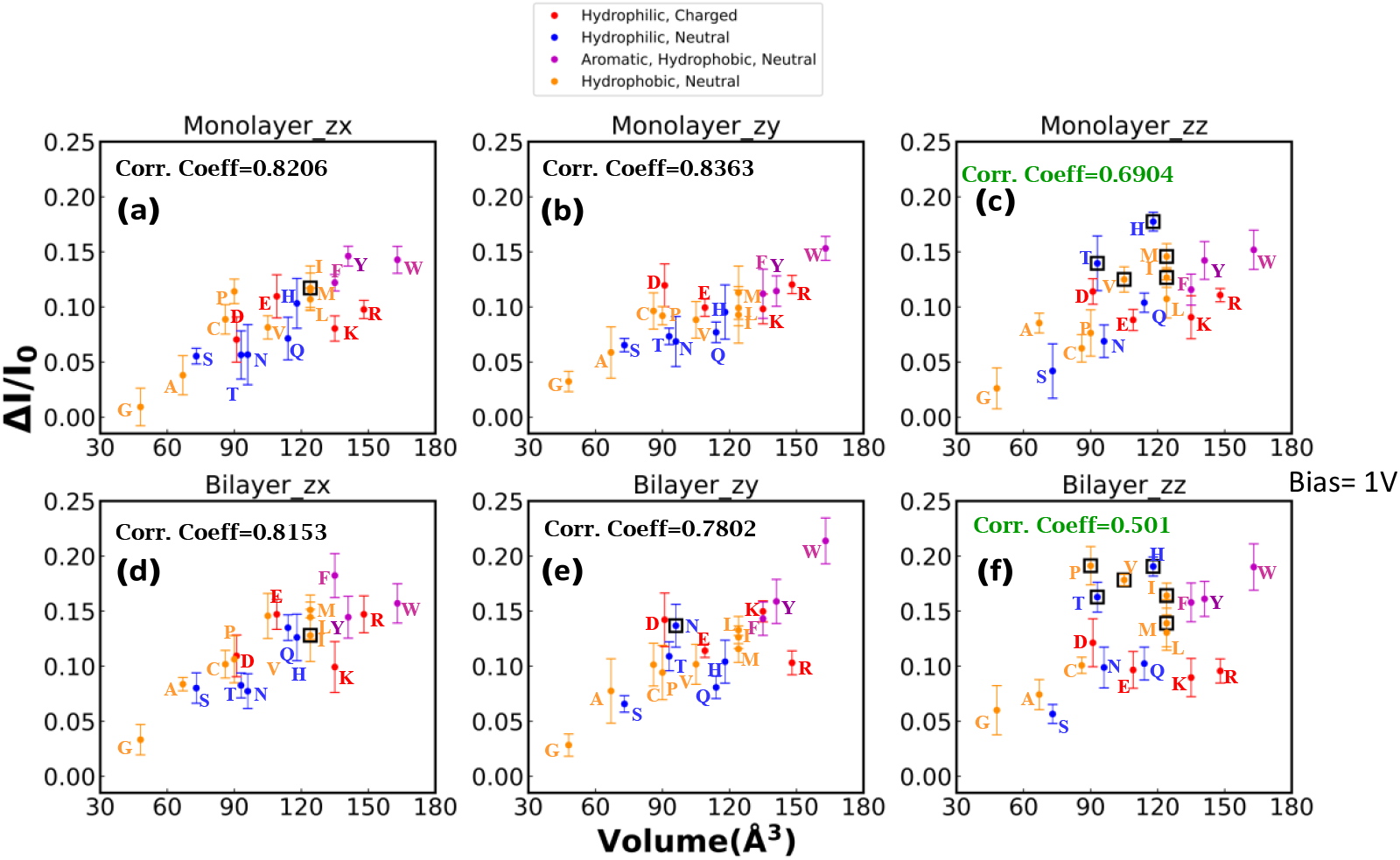
Correlation of vdW volumes and relative current blockade produced by translocating amino acids subjected to constant velocity pull. Top and bottom row represents monolayer (a, b, c) and bilayer graphene pore geometry (d, e, f), respectively. The first, second and third column represents the zx (a, d), zy (b, e), zz (c, f) orientation, respectively. A reduction of correlation can be seen in the third column (c, f), i.e., zz orientation. Error bars were produced using five replicas of the production run.

**Fig. 4.**
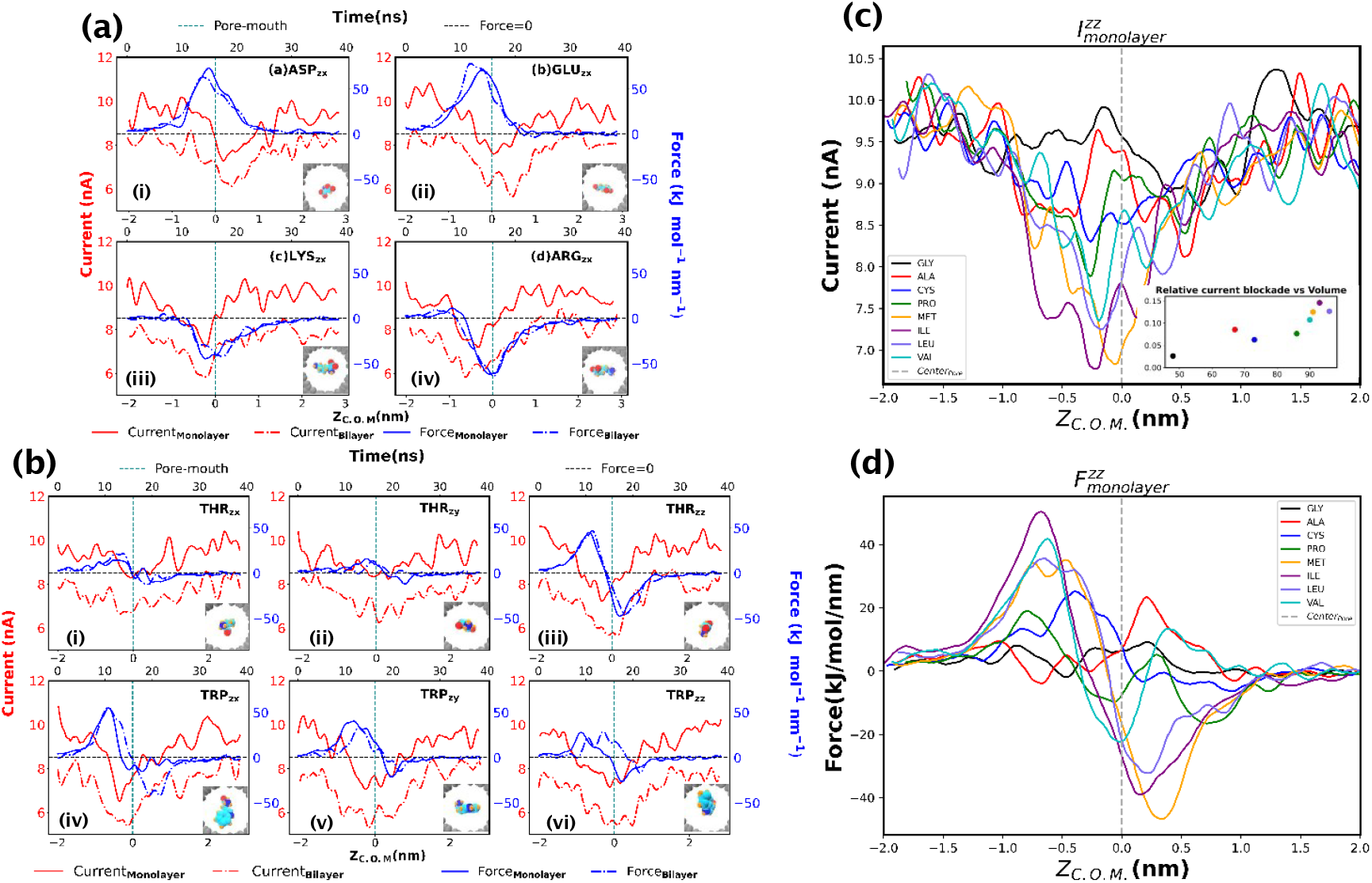
**(a)** Current and force profiles exhibiting the translocation of the charged amino acids in zx orientation have been shown here. Top row (i, ii) is for the negatively charged amino acids and we see a positive or repulsive nature of force curves. The bottom row (iii, iv) is the for the positively charged ones where we see a negative or attractive nature of force curve. The solid line represents the monolayer graphene pore geometry and the dot-dashed line is for the bilayer graphene pore geometry. We see lower value of currents for the bilayer. **(b**) Current and force profile exhibiting the translocation of the neutral amino acids THR (i, ii, iii) and TRP (iv, v, vi). The first, second and third column represents zx (i, iv), zy (ii, v), zz (iii, vi) orientation of the amino acids, respectively. The insets depict the amino acids at different orientation while translocating. Here, the force profiles contain both an attractive (-ve) and a repulsive (+ve) region, although the distinction between the regions is quite prominent for THR_*zz*,_ (iii) showing highest peak to peak difference in magnitude. Similar current dip for THR_*zz*_ and TRP_*zz*_ can be seen in the third column (iii, vi), although TRP has the highest volume among all amino acids and THR is in the mid-range. **(c)** Distinctive current Signals and **(d)** Distinctive force signals characterizing translocation of the amino acids (zz orientation) in the hydrophobic, neutral, non-aromatic category.

In zz orientation, although the aromatic group produces high relative current blockades, amino acids from relatively lower volume class (hydrophobic and non-aromatic and neutral, hydrophilic and neutral) surprisingly produce current blockades comparable to or higher than the aromatic group. The points marked with bold black squares in Fig. 3 identify these amino acids, who are responsible for decrease of correlation in zz orientation for the pulling case. These squares are also present for ASN in zy configuration (bilayer) and MET, ILE in zx. Overall, the squares are assigned for a different behavior of those amino acids. Each of these marked amino-acids in respective scenario, produce a similar, distinct shape of force-curve along with a prominent dip in the current signal, which are different from the rest of scenarios for the same amino-acid. For example, THR is marked with a square in zz for both monolayer and bilayer (Fig. 3**c, f**). If we compare the relative blockade current for THR in zz with orientation zx (Fig. 3**a, d**) or orientation zy (Fig. 3**b, e**), we see that the value is highest for zz in both the scenarios for monolayer and bilayer. In addition, the value of the relative current blockade for THR is comparable with TRP in zz orientation while for other orientations, the values are quite different. We did not expect the values to be comparable since, volume of THR and TRP are not close. If we look at Fig. 4**d**, we see that in panel **iii)**, the force curve for THR in zz, is quite distinct and higher in peak-to-peak magnitude compared to panel **i)** and **ii)**. The lowest current value in panel **iii)** is also the lowest among panel **i), ii), iii)** and both the force and current profile of THR in panel **iii)** is now similar to TRP (panel **vi**). The force profile in THR_*zz*_ which is approximately anti-symmetric about the pore, have approximately similar parts of repulsive and attractive region demonstrating a prominent approach and exit. This shape of force-curve, which one orientation produces, and other orientations do not, is not observed in case of other non-marked molecules. For TRP (Fig. 4**d i, ii, iii**), the signals are similar for all the orientation and that’s why TRP was not marked in black. The value for relative current blockade of HIS in zz for monolayer is also higher than the largest amino-acid TRP (Fig. 3**c**). We observe that while THR, VAL, HIS, MET, ILE in zz orientation exhibits this special behavior for both monolayer and bilayer, PRO in zz exhibits this only for the monolayer. This layer no. dependence exists also for MET, ILE in zx and ASN in zy. While comparing the blockades (Fig. 2 and Fig. 3) of the static and translocating amino acids, we see that the magnitude of relative current blockade for the static ones are quite higher than the value we got while pulling them. To explain this, we look at the dwell time of the amino acids at the pore. The C.O.M. of the static amino acids are anchored to the pore-center for the whole 40 ns, but the pulled amino acid stays in the region of |Z|=1nm around the pore for approximately 16 ns as per the pull rate. Previous works^6,61^ have reported a better sensing when the molecule is slowed down to reside for a longer time interval at the pore. The error bars for relative current blockades are significantly higher in case of translocating amino acids. The error bars for relative current blockades are significantly higher in case of translocating amino acids. The error-bars were calculated using the five different replicas. Trajectories for translocating amino acids have more variance compared to static ones. The amino acid interacts with the surrounding waters and ions and pore atoms more dynamically for translocation studies compared to the static cases, which leads to more fluctuation and may add to the reduction of relative current blockade during translocation. We have calculated the RMSD of the of ARG in zx orientation both for the pulled and static case and compared them in **Fig. S41** in section 13 of SI. A larger fluctuation is observed in the pulling case. The dynamics of the moving amino acids introduce the effect of confinement which also contributes to fluctuating trajectories. We have also compared overall relative current blockade variance of monolayer pore and bilayer pore in SI (**Section 8, Fig. S14**) and found the monolayer to produce lesser variance than bilayer.

### 3.3 A Closer Look into the Translocation Dynamics: Radius of Gyration and Dipole Moment of Single Amino Acids

To probe into possible origins of orientation dependence of current blockades, we calculated the radius of gyration *R*_*g*_ of the amino acids translocating through the pore.

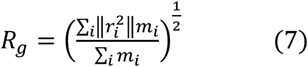

where *m*_*i*_ is the mass of atom i and *r*_*i*_ the position vector of atom i with respect to the center of mass of the molecule of interest. To provide a better understanding, we also calculated the radius of gyration about each coordinate axis (by only summing the radii components orthogonal to each axis). We are calling them projections of radius of gyration of the molecule in the plane perpendicular to the coordinate axes which are initially set to be aligned with the principal axes of the molecule. For example, projection of the radius of gyration of the molecule in XY plane (or radius of gyration about the Z axis) is given by,

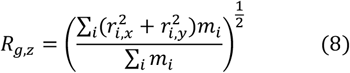

where radii components orthogonal to Z axis have participated in the summation. For each orientation of amino acid, R_g_, R_g,x_, R_g,y_, R_g,z_ have been calculated. In other words, R_g,x_, R_g,y_, R_g,z_ is indicative of the rotational tendencies of the molecule about X,Y and Z axis respectively. Fig. 5**a** shows the average R_g,z_ during the translocation of the amino acid when it is away from the pore, i.e. the average of the baseline region. In this condition, R_g,z_ and the other projections measure the shape factors associated with the molecule. We see that R_g,z_ for amino acids in zz orientation, i.e. the rotational tendency of the molecule about Z axis, unlike zx or zy, is not correlated with the vdW volume, a similar behavior we observed in the relative current blockades. This demonstrates that the orientation effect observed in Fig. 2 arises from the intrinsic way in which the shape of the amino acid scales with volume. We explain the data shown in Fig. 5**a** in the schematic Fig. 5**c**. With increasing vdW volume, the amino acid in zx or zy orientation, has longer projection along Y or X direction, which lies in the pore plane. Hence, ionic flow is hindered proportionally with the volume. In zz orientation of the amino acids, with increasing vdw volume, the amino acid has longer projection along Z axis or the perpendicular direction of the pore plane, parallel to ionic flow. Therefore, the steric resistance to ionic flow is expected to be less than the zx and zy orientation. The volume and relative current blockade correlation is thus reduced in zz orientation which can be observed in Fig. 2.

**Fig. 5.**
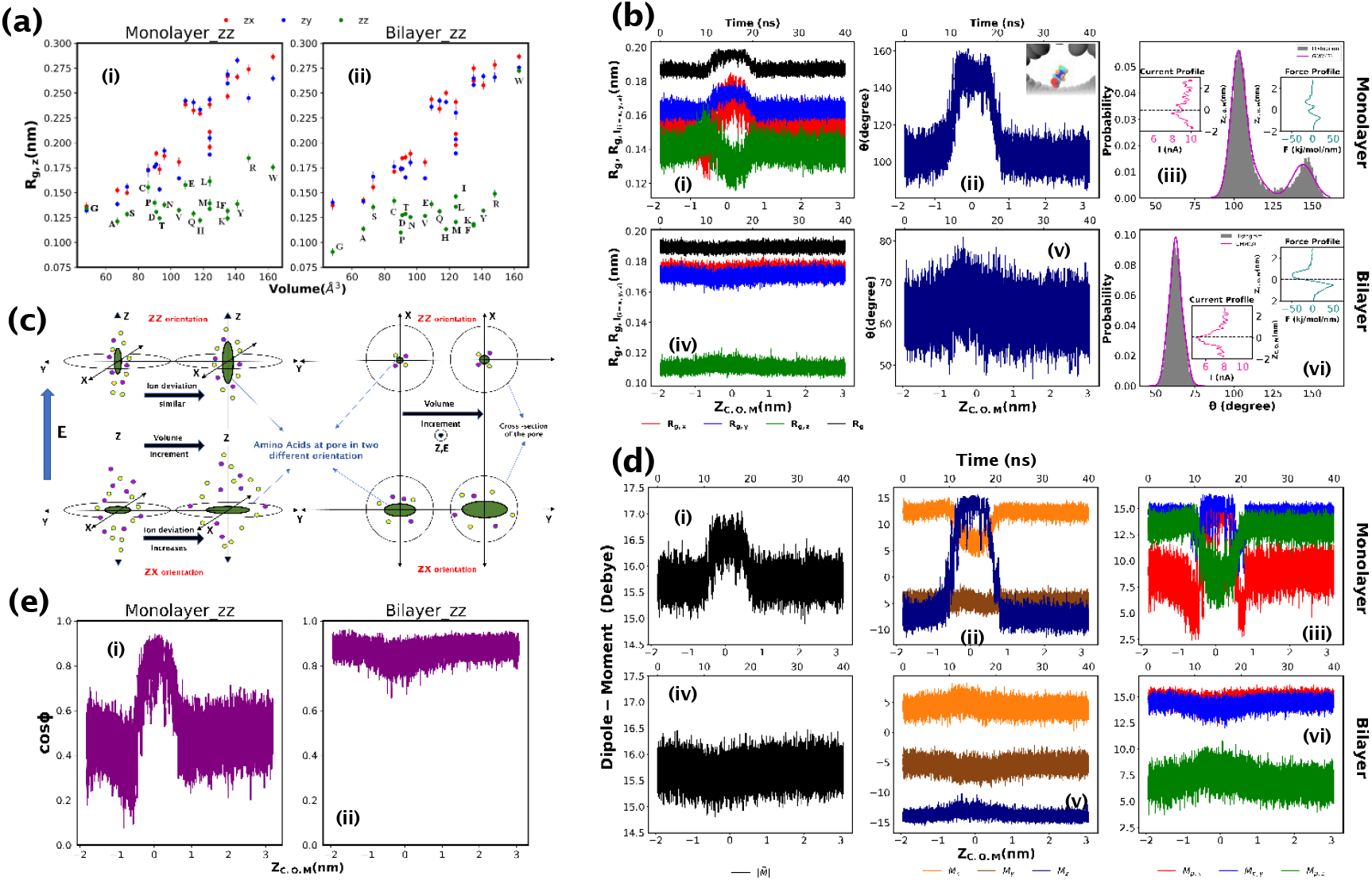
**(a)** Average R_g_ about Z axis or the projection of R_g_ in xy plane, i.e. R_g_,_z_ for each orientation of 20 amino acids when they are away from pore (baseline region) is shown here. (i) is for monolayer graphene pore geometry and (ii) is for bilayer. In both the cases we see reduction of correlation between R_g_,_z_ and the volume of the amino acids for the zz orientation of the amino acids. **(b)** First column represents the dynamic R_g_ (along with its projections) profile (i, iv). The second column (ii, v) shows fluctuation of the angle θ between the normal to the plane formed by C_α_ −C_β_ −N_backbone_ of the translocating amino acid and the Z axis. Occupation probability for θ has been plotted in the third column (iii, vi). Top (i, ii, iii) and bottom (iv, v, vi) row represents monolayer and bilayer pore geometry, respectively. Every plot is for PRO in zz orientation from the pulling studies. The insets in (iii, vi) represent the current (deep pink) and force profile (teal) in the respective setup. For bilayer pore geometry (vi), we can see higher current blockade and a prominent anti-symmetric force profile with higher magnitude as compared to monolayer (iii). It should be noticed that there is no prominent dip in the corresponding R_g_ (iv) and θ (v) profile for bilayer. The inset in (ii) shows a typical translocation of PRO for zz orientation, through monolayer graphene. **(c)** This shows the interpretation of the plot in Fig. 5a. For zx and zy orientation, the longest semi-axis of the molecule is approximately parallel with pore-plane/XY plane and for zz orientation, it is approximately perpendicular. Here we explain how increment in volume for zx and zz orientation is differently affecting the ion flow-path which possibly results in the decreased correlation in Fig. 2(c, f). **(d)** (i) and (iv) represent the dipole moment 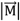 of the amino acid for monolayer and bilayer respectively. (i) shows distinct change during translocation and (iv) is comparatively flatter which is similar to b) i, iv. panels. (ii) and (v) depict different components of dipole moment vector. In (ii), M_z_ increases during translocation which also governs the total dipole moment in (i). This is also correlated with angular fluctuation in b) ii. In the last column (iii, vi), projections of the dipole moment vector have been plotted to contrast against R_g_ projections. **(e)** This describes the alignment of the dipole vector with Z axis. For monolayer graphene pore (i) we see that PRO tries to align its dipole with the Z axis during translocation. For bilayer graphene pore (ii), PRO is already aligned with dipole, as evident from the baseline. In the translocation region, the curve is flat compared to panel (i).

We show translocation profiles of Proline in Fig. 5 as an example. When we examine the dynamic profile of radius of gyration and its projections for the translocating amino acids, we notice that the profiles show a distinct jump while some of the amino acids are getting translocated (Fig. **5b.i, Fig. S15-S34**). We show (SI, section 10, **Fig. S35**) that during translocation, under confinement, in presence of electric field, the amino acids may get stretched and hence, the increase in the radius of gyration during translocation. This affects the molecule’s rotational tendency and resists its rotational diffusion. The high electric field near the pore exerts more force on the molecules, compared to when the molecule is away from the pore. It is well known in the field of nanopore research that the electrical activity is quite different around the pore compared to the bulk. The sudden potential drop across the pore gives rise to high electric field inside the pore and based on the partial charge distribution, the molecular structure responds to align the dipole moment with respect to the direction of electric-field, or here, the Z axis. This structural change is reflected in radius of gyration. Fig. 5**b** demonstrates the structural characterization of Proline or PRO during translocation in zz orientation and Fig. 6 is an example of what we mean by structural fluctuation. In monolayer pore, the force is not high (Fig. 5**b.iii**, inset with teal curve) in |*Z*_*C*.*O*.*M*._| > 1 nm region while in bilayer pore, PRO is experiencing a force of higher magnitude, (Fig. 5**b.vi**, inset with teal curve) nearly anti-symmetric about the pore (similar to THR_*zz*_ shown in Fig. 4**b**) and parallelly exhibiting an enhanced current blockade (Fig. 5**b.vi**, inset with pink curve). We have further looked into θ (Fig. 5**b (ii**,**v**)) which is the angle between the normal to the plane formed by C_α_ −C_β_ −N_backbone_ of the amino acid and the Z axis. Histogram of θ (Fig. 5**b(iii, vi**)) fitted with a gaussian mixture model was also calculated. Change in θ is reflected in the change of R_g_ and the histogram suggests that the occupation probability of θ is unimodal when the distinct force profile arises. It should be noted that having a unimodal θ distribution does not always correspond to a prominent current dip and a distinct force curve. But mostly the distribution is unimodal when that happens.

**Fig. 6.**
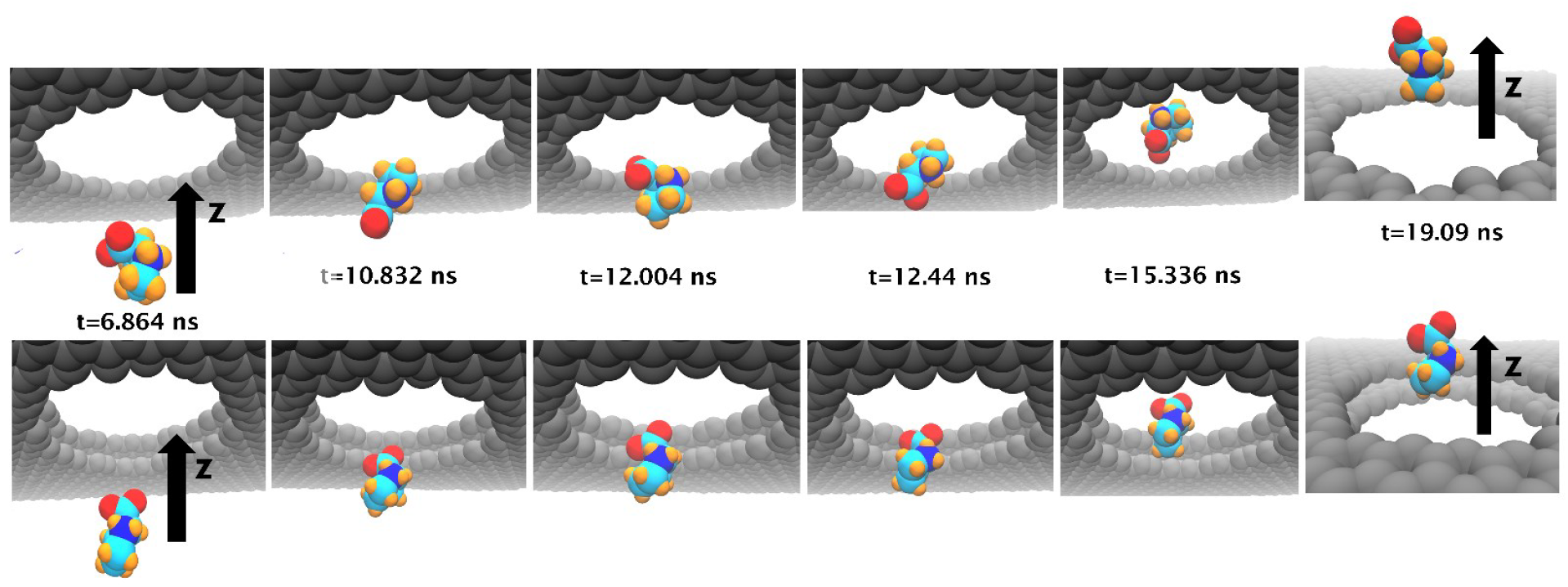
Comparison of trajectories of PRO through monolayer and bilayer graphene pore. The top row is for monolayer and the bottom is for bilayer. If we observe the two red oxygen atoms, their positions are flipping up and down in the case of monolayer. This fluctuation is absent for translocation in bilayer. The RMSD of PRO in both cases have been compared in **Fig. S40**.

Recent studies^22,23,25,27–29^ have shown the importance of dipole moment in the translocation dynamics of proteins. We now investigate how the dipole moment of the amino acids change while they translocate. We have used a non-polarizable additive force field in our simulation and the limitation of these type of force fields in dipole calculation is well known^62^. The partial charges assigned are fixed and are not adaptive to the environment. Therefore, no existence of induced dipoles via polarization. However, the high electric field in the pore region can change the intra-atomic distances of the molecule. This can cause stretching of the molecule which corresponds to the change of the value of dipole moment of the molecule or its components while the amino acid translocates through the pore. Dipole moment gives information about structural alignment, while R_g_ characterizes both structural change and volume. Still, to draw parallels with projection of R_g_, we have calculated the modulus of projection of dipole moment 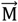 in different planes. For example, the projection in XY plane is given by

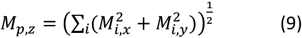

where components orthogonal to Z axis have participated in the summation. This is not the z component M_z._. For each orientation of amino acid M_p,z_ has been calculated. We find that the M_p,z_ doesn’t significantly correlate with volume in any orientation (SI section6, **Fig. S11**,**S12**). The z component of 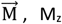 shows no prominent correlation with volume in any orientation (SI section6, **Fig. S10**) either. We characterize the dipole moment profile for PRO at zz orientation in Fig. 5**d, e**. Fig. 5**d** (**i, iv)** shows the dynamic profile of modulus of dipole moment vector of the amino acid, which correlates with panel i) and iv) of Fig. 5**b** respectively, the R_g_ profiles. In Fig. 5 **d** (ii, v), we analyze the different components of the dipole moment and Fig. 5**e** quantifies the orientation Ф of the dipole moment vector with Z axis, in terms of the cosine of Ф. We find that 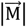 in panel **d (i, iv)** is governed by M_z_ and Ф. The changes in M_z_ and cos Ф are also correlated with θ (Fig. 5**b (ii, v**)). Fig. 5**e.i** demonstrates how the dipole alignment with the electric field (Z axis) for PRO_zz_ increases while translocating through the monolayer pore.

We will now shift our focus towards understanding the translocation behavior of the marked amino acids (Fig. 3) mentioned previously, which, mostly in zz orientation, produce a current blockade not in proportion to their volume. For Proline_zz_, this was observed in bilayer. The special nature of force curve that has been discussed earlier can be associated with the fact that in almost every case (**Fig. S20-21**,**23**,**30**,**32-34**), the R_g_ and its projection (Fig. 5**b.iv**), dipole moment of the amino acid (Fig. 5**d.iv**) and its alignment with Z axis or the electric field (Fig. 5**e.ii**), exhibits no significant change during translocation. Therefore, the dipole moment of the molecule was aligned with the electric field even when the molecule was away from pore. There is no upward jump in the R_g_ profile indicating that the molecule does not experience stretching during translocation. Consequently, the structural fluctuation of the molecule during translocation, is low (SI section 12, **Fig. S40**) compared to its other orientations. This in turn produces a better current blockade. We observe in Fig. 6 that the structural alignment of the amino acid with the translocation axis or the direction of electric field is different for monolayer and bilayer. For bilayer, where we see better current blockade, there is no structural flipping of the molecule as seen in monolayer.

### 3.4 Comparison of Relative Current Blockades with a Recent Experiment

We have compared our results with the data published by Wang et al.^46^ with single amino acids as their molecule and MoS_2_ as their membrane. From Fig. 7 we observe that relative current blockades extracted from our translocation data compare with their result in terms of following a linear trend, especially in higher volume range but not in magnitude. The relative difference between two consecutive amino acid is similar in fashion. We must keep in mind that we used graphene pore with diameter=2nm, while the experiment was done with MoS_2_ pore with diameter<0.8nm. This naturally increases the confinement effect, resulting in the mismatch between the experiment and our pulling data. A more detailed analysis can be found in SI section 4.

**Fig. 7.**
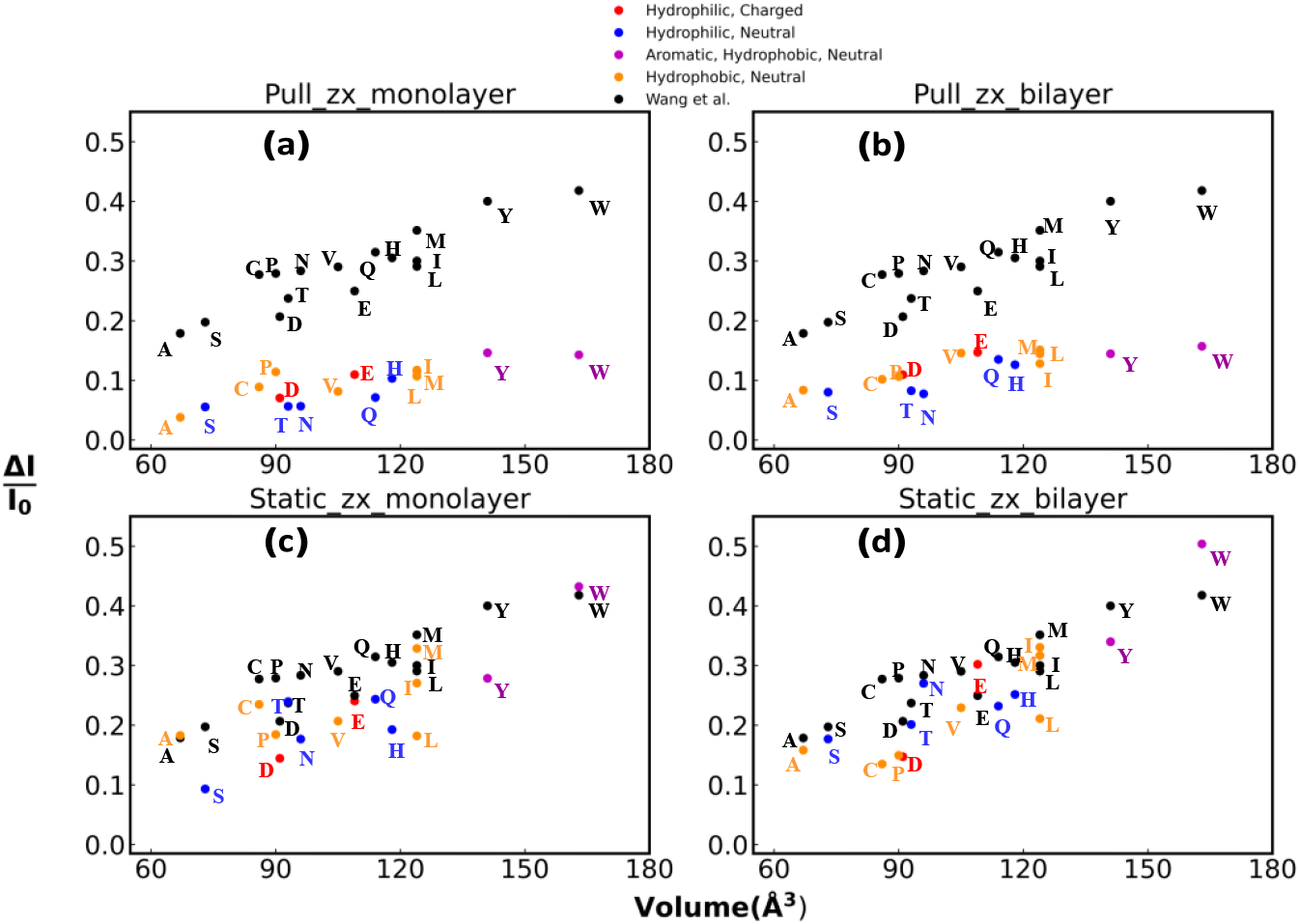
Comparison of the relative current blockades for amino acids in zx orientation, with the experiment by Wang et al^46^. The first row (a, b) is where we compare the data for amino acids being pulled through the pore. We see relative difference between two consecutive amino acids are approximately similar in most cases. In the second row (c, d) we compare the data for amino acids kept static at the center of the pore mouth. The values of the relative current blockades are now comparable with the experiments for a greater number of amino acids. The first (a, c) and second (b, e) column represents monolayer and bilayer graphene pore geometry, respectively.

When we contrast our data for the static amino acids against the experiment, we observe in Fig. 7 that the magnitude of the relative current blockades for a greater number of amino acids are now comparable with the experimental value. To explain this, we compare the residence time of the molecule at the pore. Dwell time at the pore is 2.5 times larger for the amino acids static at the pore compared to the case where amino acids are being pulled through the pore. The former is more proximal to the translocation time scales of the experiment. Here, comparison is shown for zx orientation only, the rest can be found in SI, section 4, **Fig. S5, S7**.

## 4. Conclusions

This study was designed to explore the nature of single amino acid translocations through graphene nanopores and to probe the effect of orientation at single amino acid level. We successfully characterize the translocation of single amino acid for all the twenty amino acids and demonstrate distinctive current and force profiles. We find that the relative current blockade is dependent not only on the volume, but also the orientation of the amino acid at the pore. This orientation effect is stronger when the amino acid is static at the pore compared to when the amino acid is translocating across the pore. We argue that this is due to the increased dwell time of the amino acid at the pore which also improves relative current blockade magnitudes. The translocating amino acids are also subjected to more fluctuation arising out of the environment, adding to the cause of diminished orientation effect in pulling data. To get a microscopic understanding of the orientation effect we calculated radius of gyration (R_g_) and its projections in three orthogonal planes. We found reduction of correlation between volume and projection of radius of gyration in XY plane in zz orientation of the amino acids even when the molecule is away from sensing region of pore. This shows that intrinsic structural effect is involved in the prominent reduction of correlation between volume and relative current blockade for static amino acids. In other words, the loss of correlation arises from the way the amino acid in zz orientation, with increasing volume, creates a longer projection in a direction perpendicular to the pore plane and parallel to the direction of ionic flow. In pulling studies, we find some amino acids, mostly in zz orientation, produce relative current blockade not in proportion with their volume. We see these amino acids experience force of higher magnitude and distinct shape while translocating through the pore. After investigating the dynamic profiles of radius of gyration and dipole moment, we conclude that this high force is to resist structural fluctuation of those amino acids during translocation since their dipole moment were already aligned with the electric field even when they were away from the pore. These amino acids show no jump in the R_g_ profile indicating no stretching during translocation adding to the cause of lower structural fluctuation. The lesser the fluctuation while translocating, the better the current blockade. Several questions remained unanswered such as how the chemical compositions of the amino acids affect the current blockade or why certain amino acids behave differently with different number of graphene layers. These are the fundamental aspects we are interested to address in the future. We also want to add different neighbors to the amino acid and gradually build up a chain to explore the neighbor effects in protein translocations as an extension of this work.

## Data availability

The data supporting this article have been included as part of the Supplementary Information and are available at https://doi.org/10.5281/zenodo.14793858.

## Acknowledgements

This work was supported by the Scientific and Useful Profound Research Advancement (SUPRA) Program of the Science Engineering Research Board (SERB) under Grant SPR/2021/000275. P.S. acknowledges the PhD fellowship support from DST, India. A.U. thanks Dr. Subhadeep Basu for fruitful discussions.

