## Supplementary Information for "Orientation Dependence of Current Blockade in Single Amino Acid Translocation through a Graphene Nanopore^†^"

### Contents


1 Simulation Runtime 4

2 Pore Characteristics 5

3 Processing of raw current and force data 6

4 Comparison of relative current blockades with Experiment 7

5 Comparison of Average $R_{g,z}$ between In (Translocation) and Out(Baseline) of the Sensing Region of the Pore 10

6 Volume correlation with the Dipole Moment 11

7 Force correlation with Volume 13

8 Variance of Relative current blockade: Comparison of Layer Numbers 14

9 Pulling Profiles 15

10 Stretching of the amino acid during translocation 36

11 Categorical comparison of Current and Force profile 38

12 RMSD of $\mathrm{PRO}_{zz}$ during translocation. 40

13 RMSD of the amino acid, comparison of fixed with pulled: Example with $\mathrm{ARG}_{zx}$ 41

14 Influence of pore-passivation and adding charge to the membrane on orientation effect 42

15 Number of atoms and molecules in the simulations. 45

16 The probability density of currents for different amino acids. 45

17 Structures and Properties of the Twenty Amino Acids 46

### 1 Simulation Runtime

| **Studies** | **NVT equilibration** | **NPT equilibration** | **NVT production** | **Replicas** | **Runtime** |
|---|---|---|---|---|---|
| **Openpore** | 2 ns | 2 ns | 10 ns | No replica, but 13 different Electric fields for each monolayer and bilayer | (2+2+10)×11×2=**308 ns** |
| **Static amino acid** | 5 ns | 5 ns | 40 ns | 5 replicas for each orientation zx,zy,zz for each monolayer and bilayer | 5+5+(40×5×3×2)=**1210 ns** |
| **Pulling** | 5 ns | 5 ns | 40 ns | 5 replicas for each orientation zx,zy,zz for each monolayer and bilayer | 5+5+(40×5×3×2)=**1210 ns** |
| **Cumulative runtime** | −− | −− | −− | −− | 2728 ns = **2.728** $\mu$**s** |

Table S1

#### 2 Pore Characteristics

The I-V characteristics of the pores are depicted in Fig(2). Even when the voltage is zero, the current through monolayer pore is slightly negative (inset of Fig.S1) which is indicative of thermal motion of ions. In case of bilayer, the zero bias current is almost zero which shows that the mobility of ions have been reduced compared to monolayer. The pore conductance was calculated from the slope of the linear fit. $\sigma_{bilayer}$=8.27 nS, $\sigma_{monolayer}$=10.41 nS which lies in the neighborhood of the previously reported values[1–3]. The openpore current through the bilayer pore is lower in magnitude compared to monolayer pore. This has also been reflected in the current profile involving amino acid translocation, where the current through the bilayer graphene pore was consistently lower than the monolayer pore.

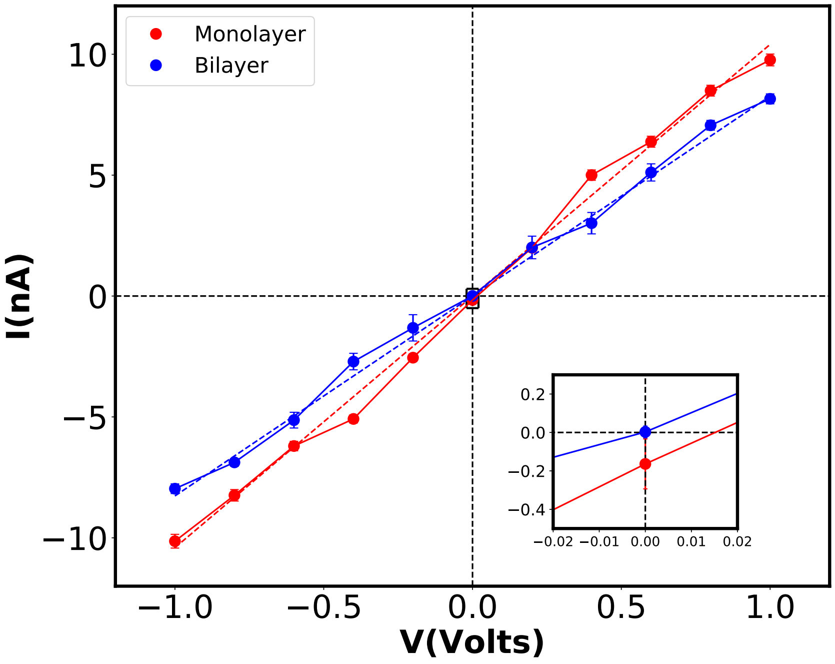


Fig. S1: IV characteristics of the pores

In the following, we show the open-pore current trace for monolayer graphene at 1V bias.

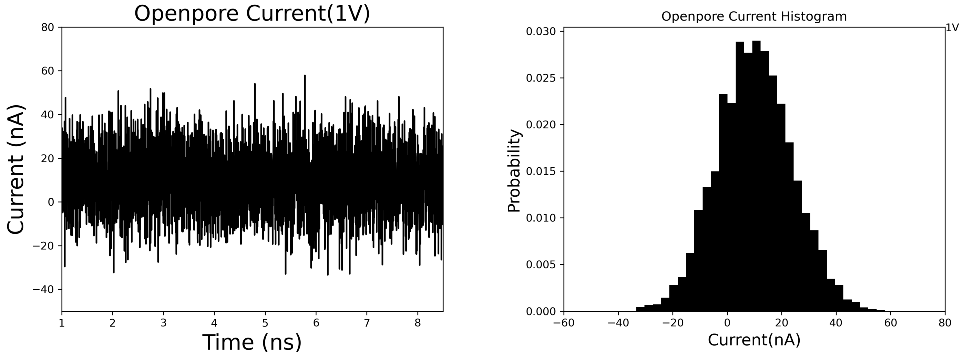


Fig. S2: Open-pore currrent trace and its histogram

### 3 Processing of raw current and force data

Here we provide a comparison of signal before and after the processing we described in the "Method" section of our manuscript.

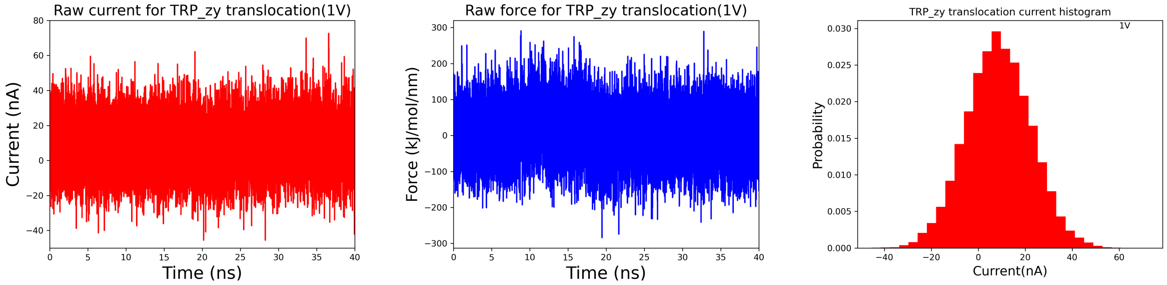


Fig. S3: A typical replica of Raw Current and Force signal along with histogram of the current trace (TRP$_{zy}$ translocation)

After processing, we average the processed trace of each replica to obtain the final curves plotted in red and blue, in the following Fig.S4

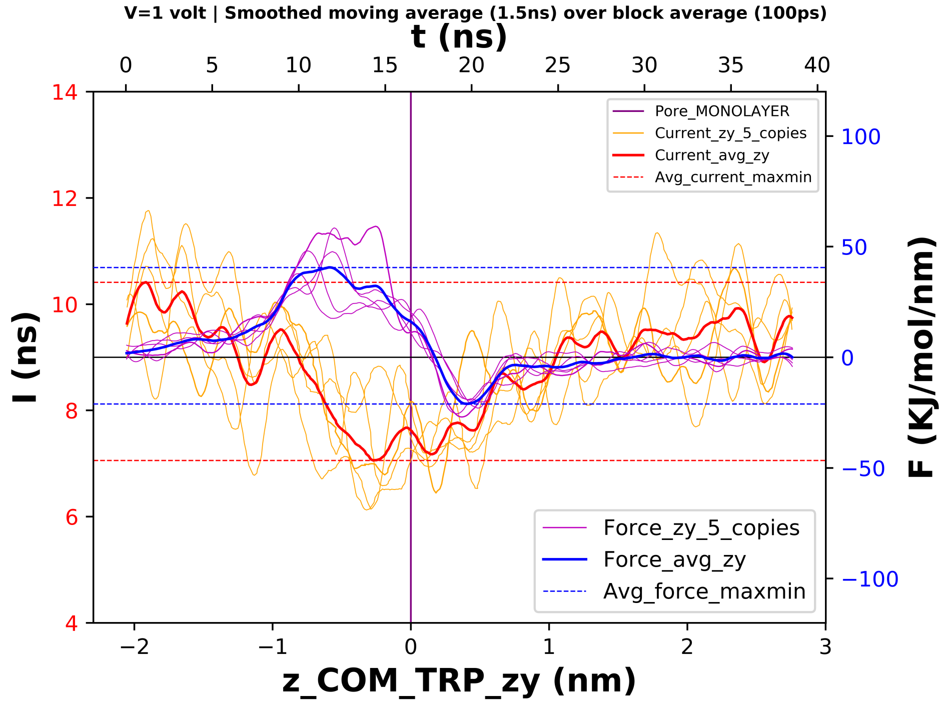


Fig. S4: Processed trace

### 4 Comparison of relative current blockades with Experiment

Here we show the comparison of our relative current blockades with the experimental data[4] for all orientations. We also provide plots with relative errors, calculated using $\frac{|Exp.Data - Sim.Data|}{Exp.Data}$ as a comparison metric. The relative error in every orientation rarely goes below 0.5 for pulling and rarely goes beyond 0.5 for the static case. In Fig.S6, the static case, the relative errors are similar for monolayer and bilayer. In Fig.S8, the pulling case, the relative errors improve for bilayer and the zz orientation has the most number of points below 0.5, as evident from the spread of the points. The graphene bilayer thickness is also more comparable to the $MoS_2$ membrane used in the experiment. The pore diameter for device 15 used in the referred work is less than 0.8nm. This naturally increases the confinement effect, resulting in the mismatch with our pull data. Also, in the experiment, the optimized range of voltage bias was 150-200mV, where we have used 1V bias. In addition to the fact that our membranes are different, if we compare our bilayer graphene open-pore current with their $MoS_2$ open-pore current, we see ours is 1.82 nA (D=2nm) at 250 mV where theirs is approximately 0.8 nA (D<0.8nm) at 200mV, which seems reasonable.

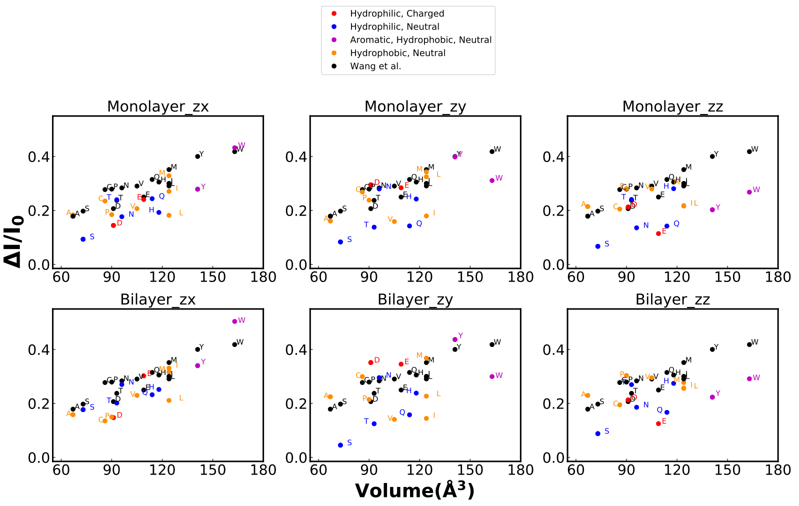


Fig. S5: Comparison with the experiment: Static

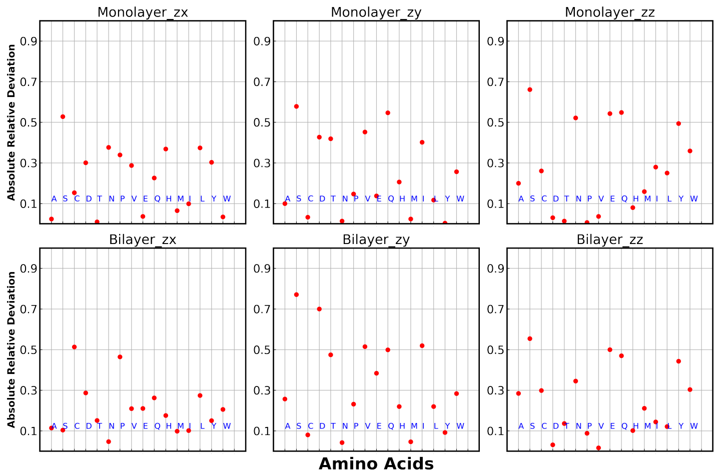


Fig. S6: Deviation relative to the experiment: Static

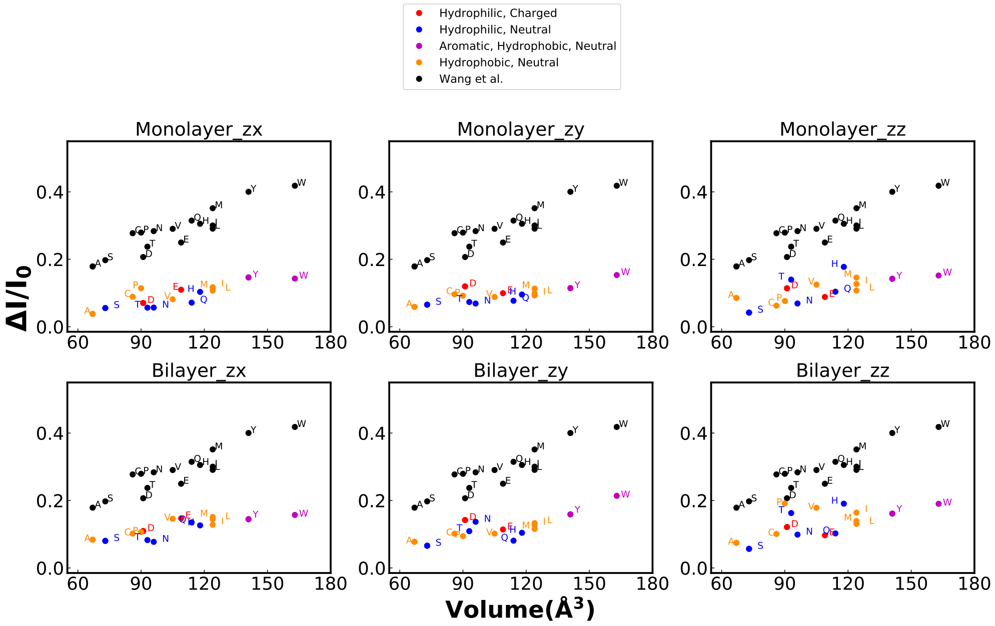


Fig. S7: Comparison with experiment: Pulling

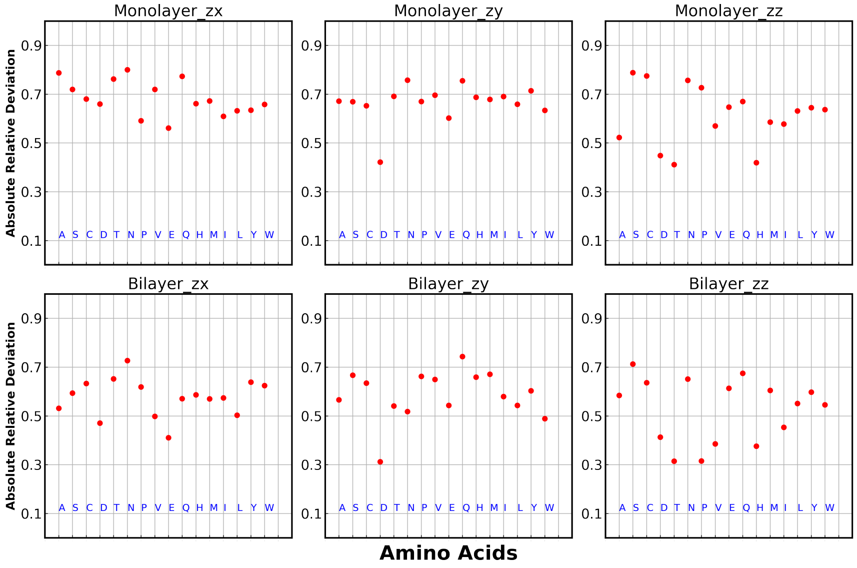


Fig. S8: Deviation relative to experiment: Pulling

### 5 Comparison of Average $R_{g,z}$ between In (Translocation) and Out(Baseline) of the Sensing Region of the Pore

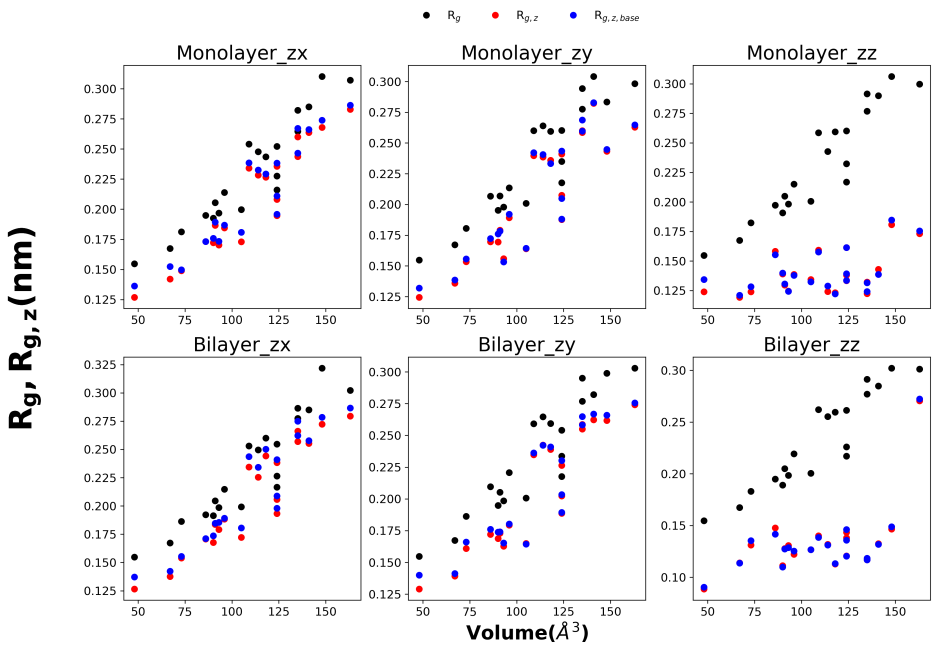


Fig. S9: $R_{g,z}$ in and out of $|Z_{C.O.M.}| = 1nm$ and overall $R_g$: Pulled Amino Acids

We can't see much difference when the molecule is in proximity or away from pore for $R_{g,z}$, as per Fig.S9. The total $R_g$ does not show any decrease in value as opposed to $R_{g,z}$ for zz orientation, which is why examination of $R_g$ about different axes (the projections of $R_g$ in different planes) was crucial.

### 6 Volume correlation with the Dipole Moment

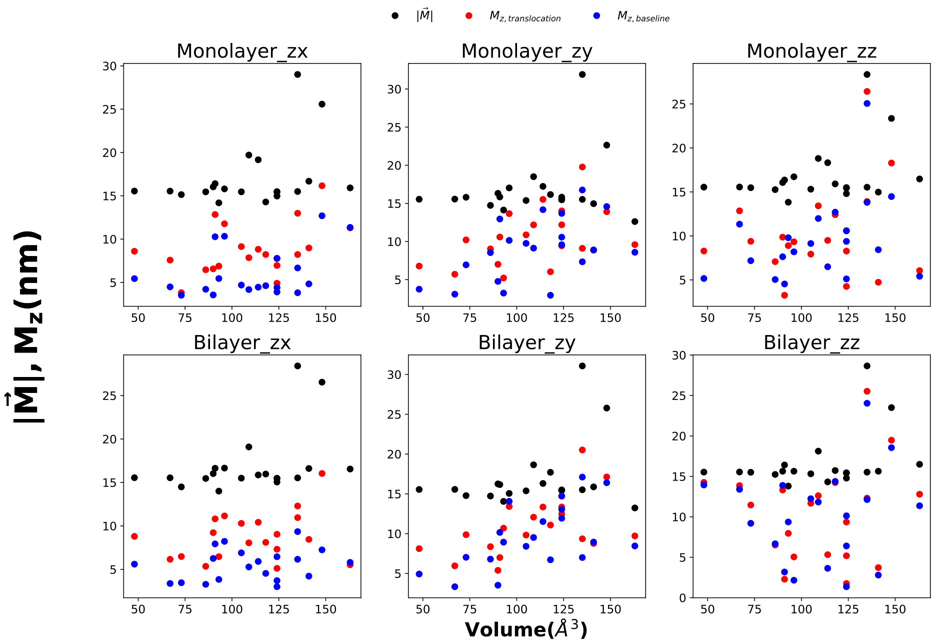


Fig. S10: $M_z$ in and out of $|Z_{C.O.M.}| = 1nm$ and overall $|\vec{M}|$: Pulled Amino Acids

We can't observe any orientation specific dependence of the volume correlation of the magnitude of total dipole moment or its z component (10). They are poorly correlated with volume.

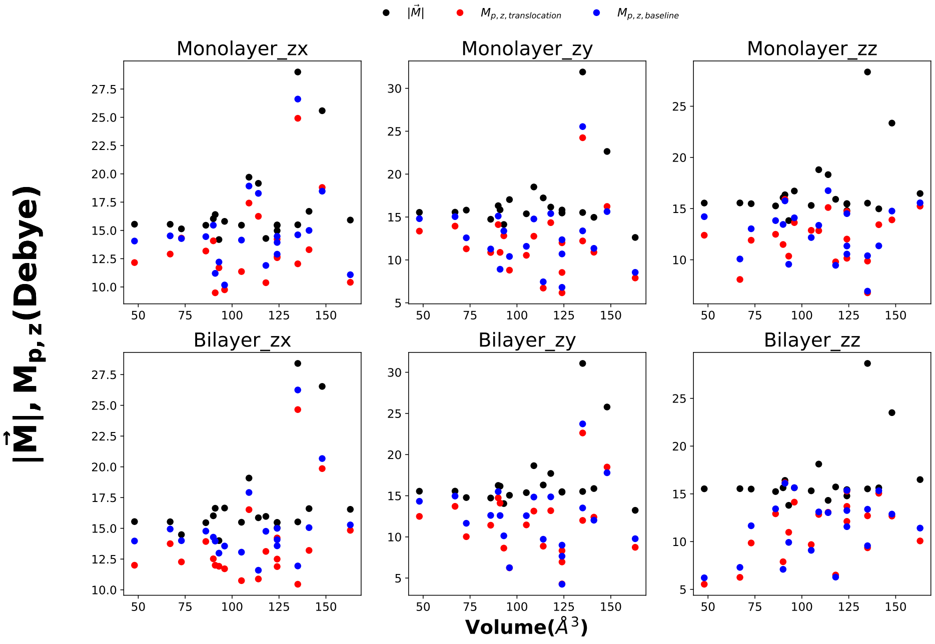


Fig. S11: $M_{p,z}$ in and out of $|Z_{C.O.M.}| = 1nm$ and overall $|\vec{M}|$: Pulled Amino Acids

We have investigated the projection $M_{p,z}$ of the dipole moment vector of the amino acids in the pore plane, to correlate with the rotational tendencies of the amino acids about the Z axis $R_{g,z}$ or the projections of $R_g$ in the pore plane. In Fig.S11 and Fig.S12 we show that $M_{p,z}$ or the projection of dipole moment in the pore-plane is not correlated with volume.

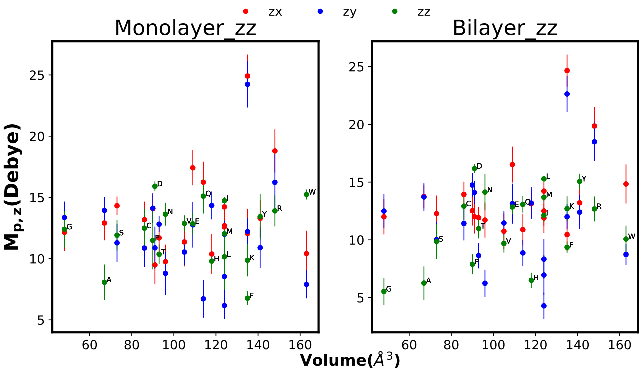


Fig. S12: $M_{p,z}$ in $|Z_{C.O.M.}| = 1nm$ for different orientations: Pulled Amino Acids

#### 7 Force correlation with Volume

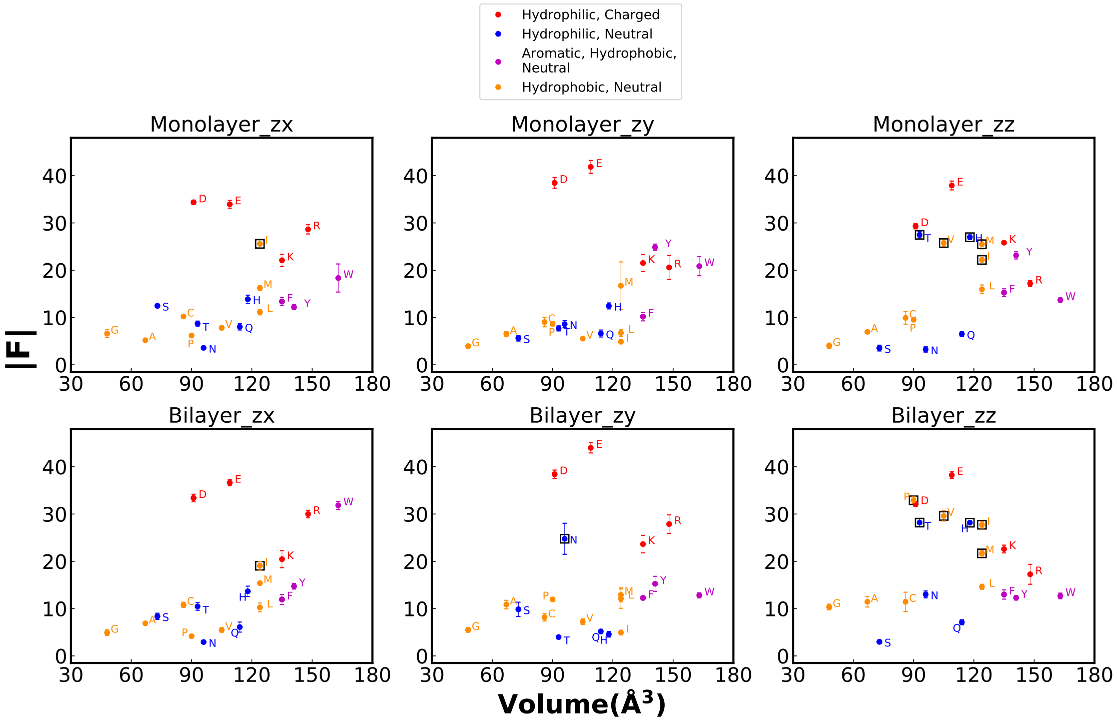


Fig. S13: Correlation of vdW volumes and force felt by amino acids while passing through the pore during pulling.

We see that the force felt by the amino acid in the vicinity of the pore is not well correlated with the volume of it. ASP, GLU which are negatively charged amino acids, are pretty distinguishable in terms of force as was reported previously for homopeptides[5,6]. The lack of correlation in zx and zy is due the fact that mostly, charged class of amino acids are located far away from the rest of others in terms of force. But for zz, the amino acids marked with squares join in. It also might be noticed that, out of six cases (3 orientations $\times$ 2 layers), five have a member of the neutral, hydrophilic group as a lowest member in the force axis.

### 8 Variance of Relative current blockade: Comparison of Layer Numbers

In this section, we want to compare variance in relative current blockade produced by monolayer and bilayer pore. For that we have gathered relative current-blockade for each run for each orientation. So each amino acid will have 5(replica) ×3(orientation)=15 data points for each of the static and pulling scenario. The variance for each amino acid was calculated from these 15 data points. In the figure Fig.S14, we show the

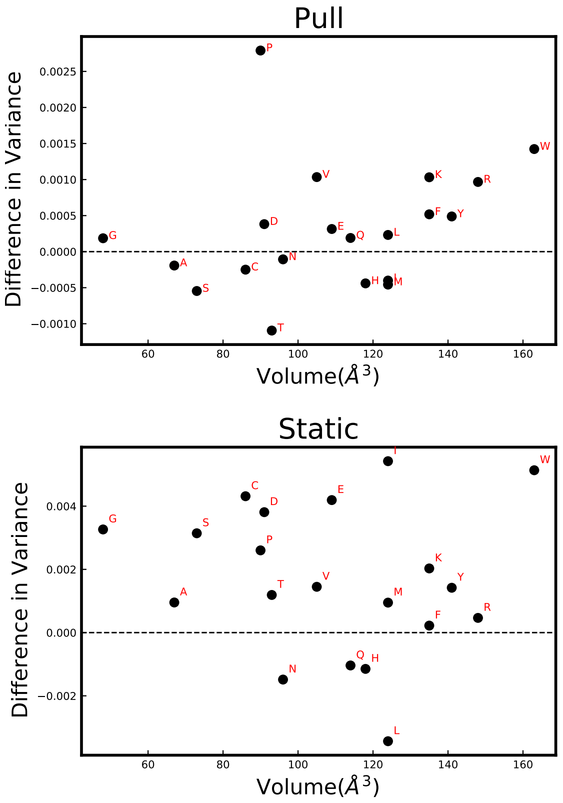


Fig. S14: Variance difference in Relative Current Blockades of Bilayer pore w.r.t. Monolayer pore. Top pannel is for the pull studies and bottom one is for the static case.

difference of bilayer pore with respect to the monolayer pore. In both the pull and static case, most amino acids lie above the zero line. This shows that the bilayer pore variance is usually higher than that of the monolayer.

### 9 Pulling Profiles

In this section, we gather all the pulling profiles for each of the amino acids.

The charged, hydrophilic class consists of ASP Fig. S15, GLUFig. S16, LYSFig. S17,ARGFig. S18.

The neutral, hydrophilic class consists of SERFig. S19, THRFig. S20, ASNFig. S21, GLNFig. S22, HISFig. S23.

The aromatic, hydrophobic, neutral class consists of PHEFig. S24, TYRFig. S25, TRPFig. S26.

The Non-aromatic, hydrophobic, neutral class consists of GLYFig. S27, ALAFig. S28, CYSFig. S29, PROFig. S30, LEUFig. S31, ILEFig. S32, METFig. S33, VALFig. S34.

We plot the translocation data for each of the twenty amino acids in twenty individual pages. In each of the following pages in this section, the first (1st row) subfigure shows the current and force profiles for different orientations. In the next row, we show the angle $\theta$ we measured as a metric of structural change (the angle between the normal to the plane formed by $C_\alpha - C_\beta - N_{backbone}$ of the amino acid and the Z axis, for GLY, the line joining $C_\alpha - N_{backbone}$ is considered instead of the plane as there is no $C_\beta$ ) in navy blue (2nd row,left) and corresponding angle occupation probability in gray plots(2nd row,right). In the following row, we demonstrate the R$_g$ profiles(3rd row,left) and the profile for dipole moment and their components(3rd row, right). In the last row we provide plots showing how the alignment of the dipole moment vector $\vec{M}$ with the Z axis varies. The cosine of the angle between Z and $\vec{M}$, $\phi$ has been plotted in this sub-figure. $\theta$, M$_z$ both were plotted with navy blue to show their correlation. From their profiles, we notice they are also correlated with cos$\phi$, plotted in purple as the last subfigure.

Fig. S15

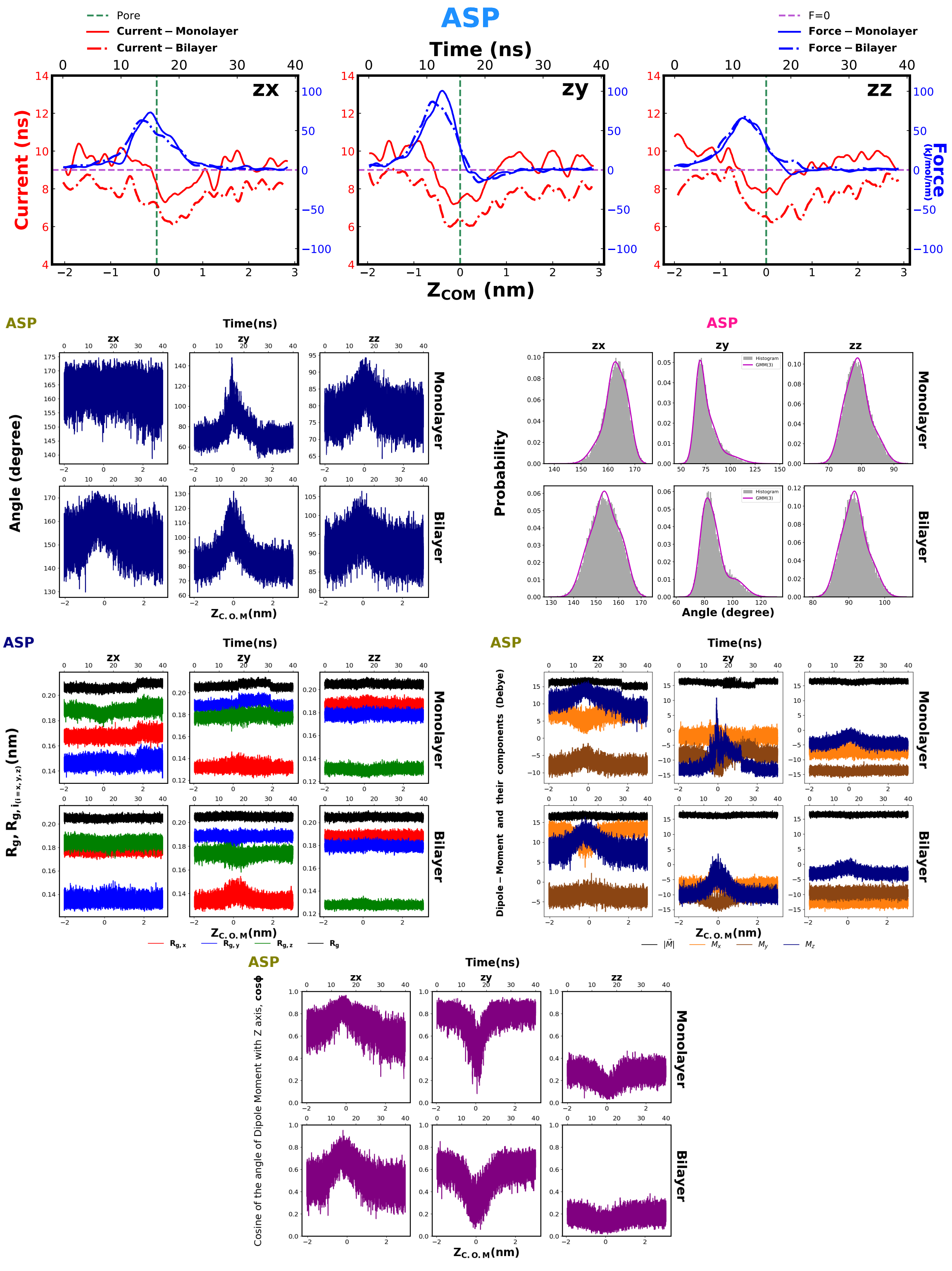
ASP
Pore
Current – Monolayer
Current – Bilayer
F=0
Force – Monolayer
Force – Bilayer
Time (ns)
zx
zy
zz
Current (ns)
Force (kJ/mol/nm)
$Z_{COM}$ (nm)
Time(ns)
Angle (degree)
Monolayer
Bilayer
$Z_{C.O.M}$(nm)
Probability
Histogram
GMM(3)
Angle (degree)
$R_g$, $R_{g,i_{(i=x,y,z)}}$(nm)
$R_{g,x}$
$R_{g,y}$
$R_{g,z}$
$R_g$
Dipole – Moment and their components (Debye)
$|\vec{M}|$
$M_x$
$M_y$
$M_z$
Cosine of the angle of Dipole Moment with Z axis, cosϕ

Fig. S16

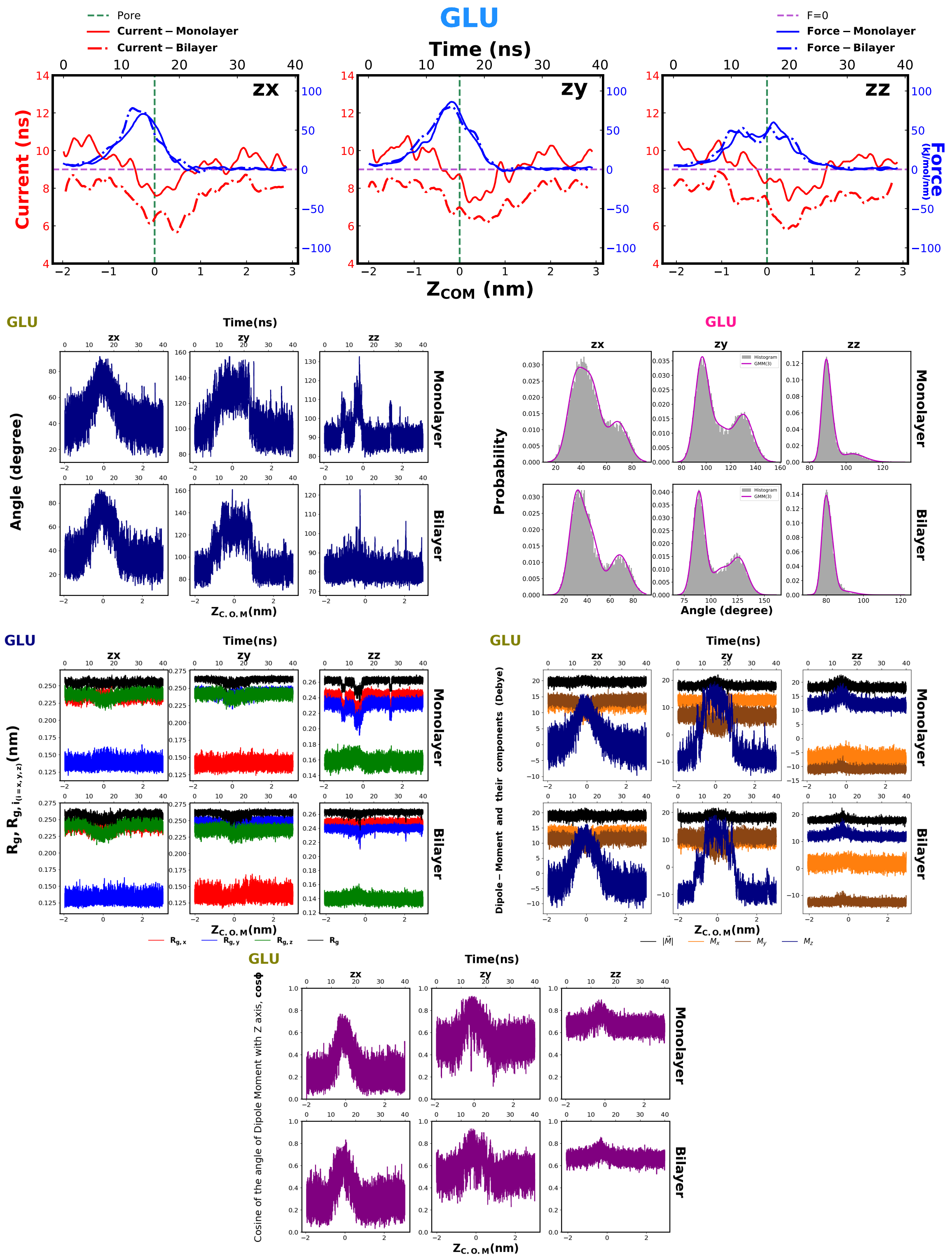
GLU
Pore
Current – Monolayer
Current – Bilayer
F=0
Force – Monolayer
Force – Bilayer
Time (ns)
zx
zy
zz
Current (ns)
Force (kJ/mol/nm)
ZCOM (nm)
Angle (degree)
Monolayer
Bilayer
Time(ns)
Zc.o.M(nm)
Probability
Histogram
GMM(3)
Rg, Rg, i(i=x,y,z)(nm)
Rg,x
Rg,y
Rg,z
Rg
Dipole – Moment and their components (Debye)
|M|
Mx
My
Mz
Cosine of the angle of Dipole Moment with Z axis, cosϕ

Fig. S17

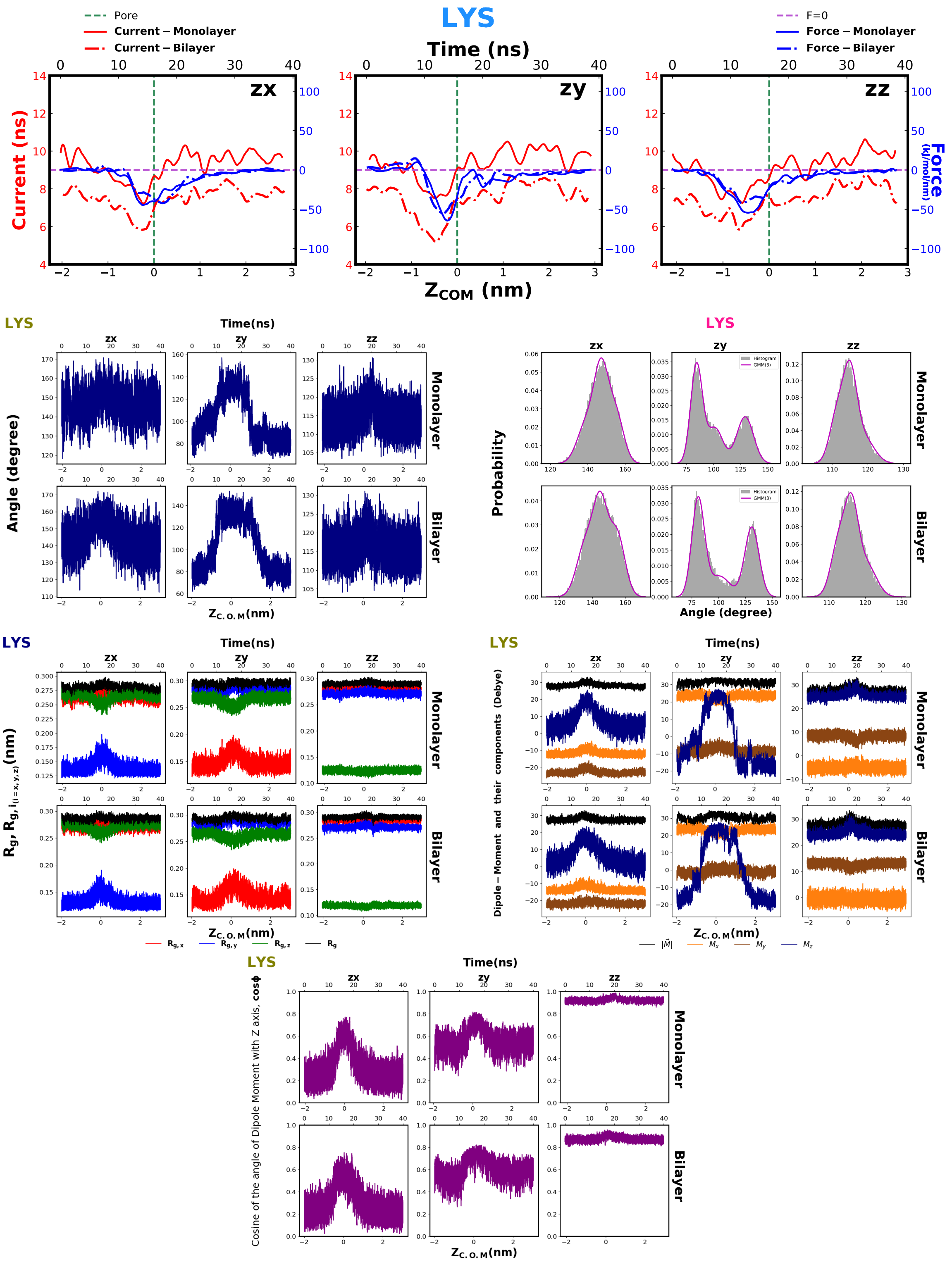

LYS
Pore
Current – Monolayer
Current – Bilayer
F=0
Force – Monolayer
Force – Bilayer
Time (ns)
zx
zy
zz
Current (ns)
Force (kJ/mol/nm)
$Z_{COM}$ (nm)
Angle (degree)
Monolayer
Bilayer
Time(ns)
$Z_{C.O.M}$(nm)
Probability
Histogram
GMM(3)
Angle (degree)
$R_g$, $R_{g,i(i=x,y,z)}$(nm)
$R_{g,x}$
$R_{g,y}$
$R_{g,z}$
$R_g$
Dipole – Moment and their components (Debye)
$|\vec{M}|$
$M_x$
$M_y$
$M_z$
Cosine of the angle of Dipole Moment with Z axis, cosϕ

Fig. S18

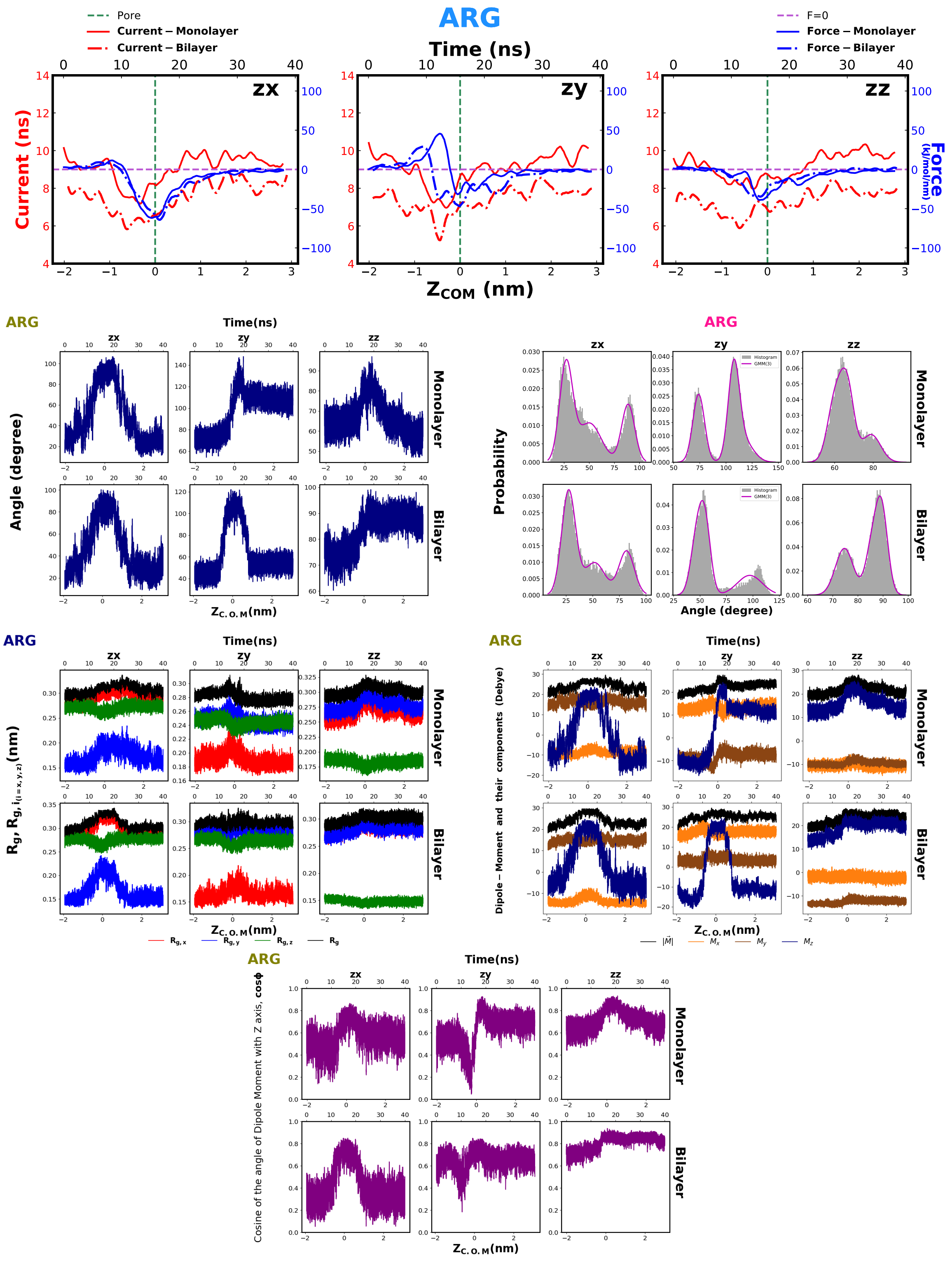

ARG
Pore
Current – Monolayer
Current – Bilayer
F=0
Force – Monolayer
Force – Bilayer
Time (ns)
zx
zy
zz
Current (ns)
Force (kJ/mol/nm)
Z_COM (nm)
ARG
Time(ns)
Angle (degree)
Monolayer
Bilayer
Z_C.O.M(nm)
ARG
Probability
Histogram
GMM(3)
Angle (degree)
ARG
Time(ns)
Rg, Rg, i(i=x,y,z)(nm)
Rg,x
Rg,y
Rg,z
Rg
Z_C.O.M(nm)
ARG
Time(ns)
Dipole – Moment and their components (Debye)
|M|
Mx
My
Mz
Z_C.O.M(nm)
ARG
Time(ns)
Cosine of the angle of Dipole Moment with Z axis, cosϕ
Z_C.O.M(nm)

Fig. S19

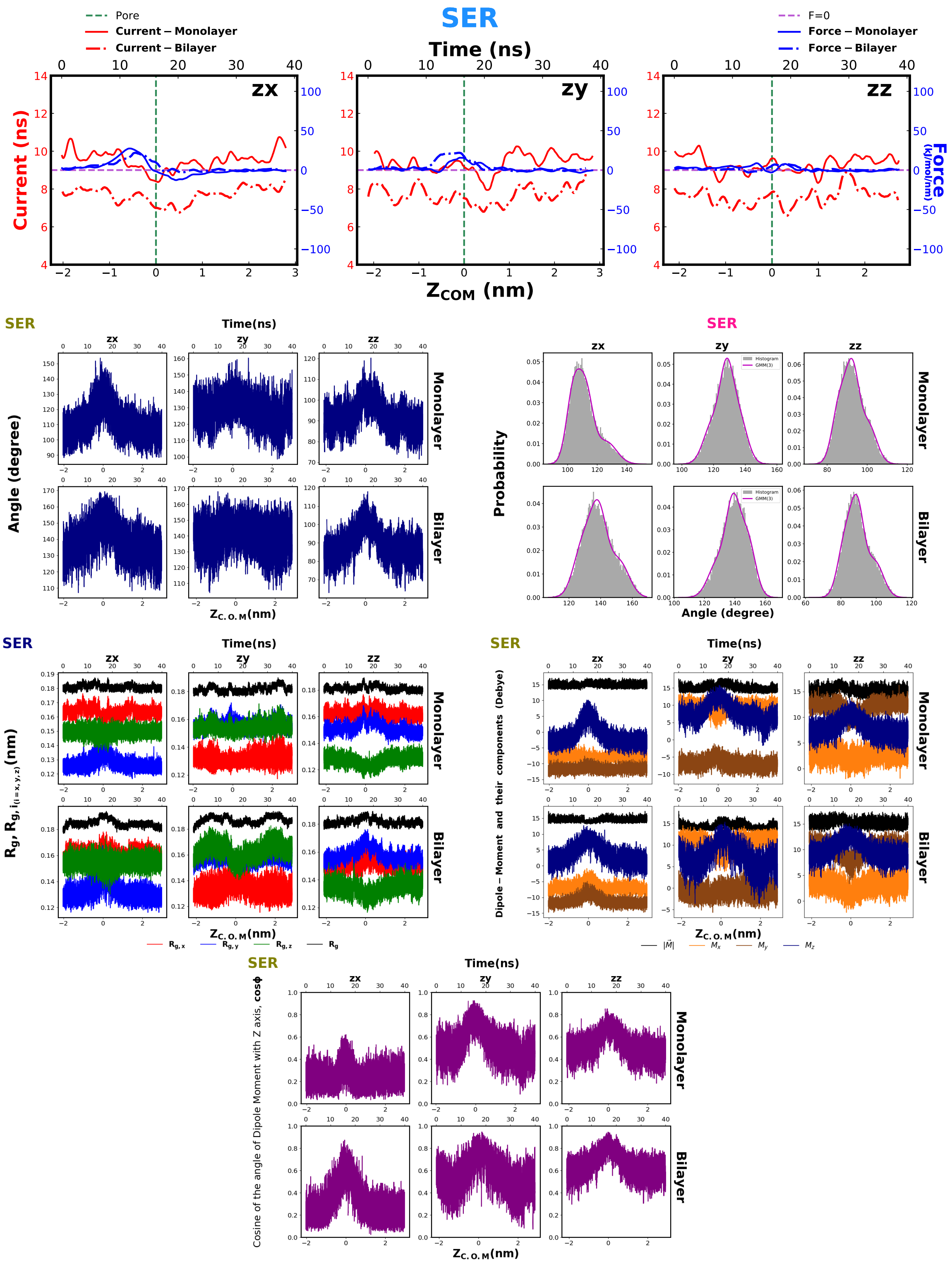

SER
Time (ns)
Pore
Current – Monolayer
Current – Bilayer
F=0
Force – Monolayer
Force – Bilayer
zx
zy
zz
Current (ns)
Force (kJ/mol/nm)
$Z_{COM}$ (nm)
Angle (degree)
Monolayer
Bilayer
Probability
Histogram
GMM(3)
$Z_{C.O.M}$(nm)
$R_g$, $R_{g,i_{(i=x,y,z)}}$(nm)
$R_{g,x}$
$R_{g,y}$
$R_{g,z}$
$R_g$
Dipole – Moment and their components (Debye)
$|\vec{M}|$
$M_x$
$M_y$
$M_z$
Cosine of the angle of Dipole Moment with Z axis, cosϕ

Fig. S20

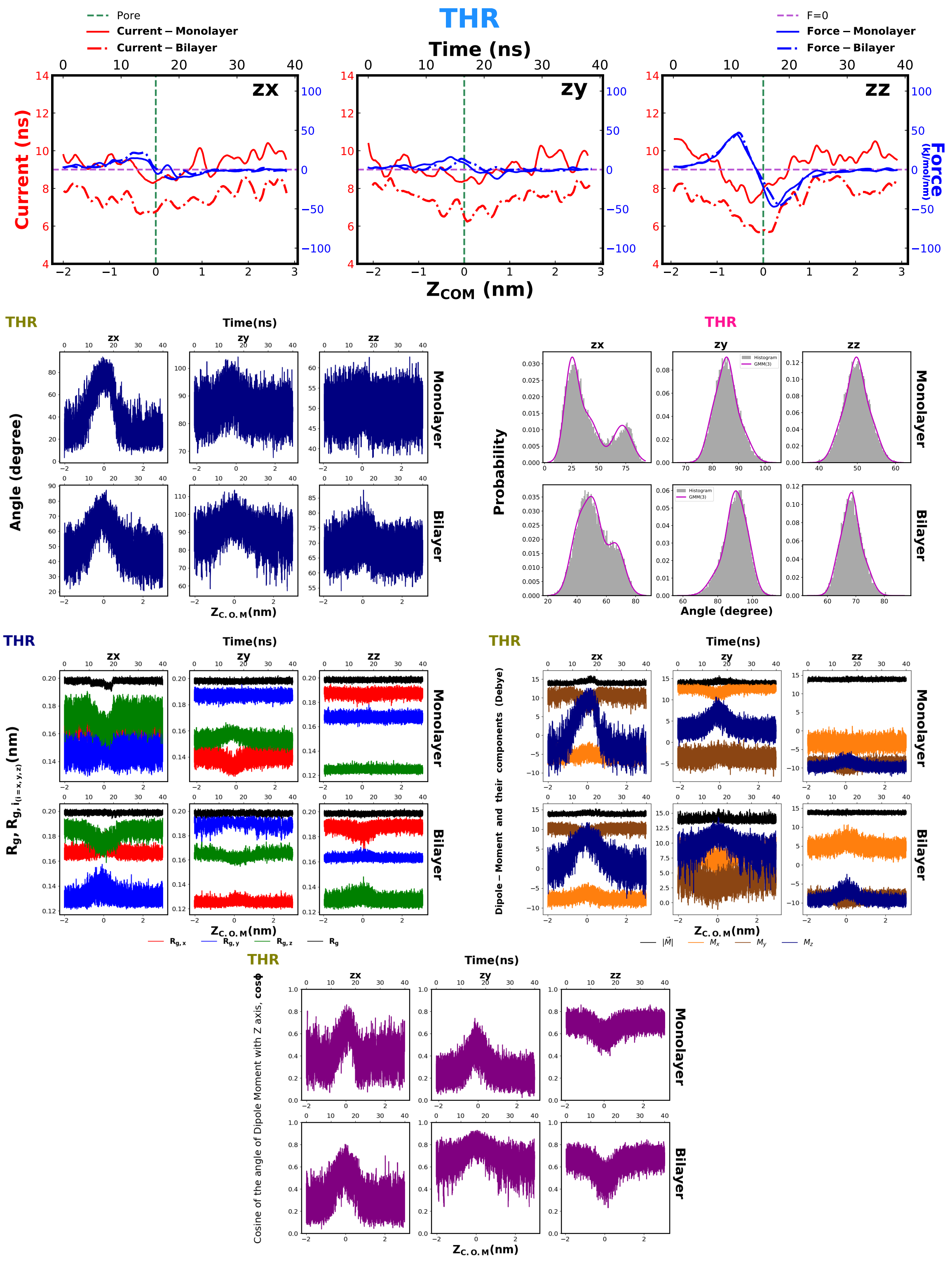
THR
Pore
Current – Monolayer
Current – Bilayer
F=0
Force – Monolayer
Force – Bilayer
Time (ns)
zx
zy
zz
Current (ns)
Force (kJ/mol/nm)
Z_COM (nm)
Angle (degree)
Probability
Histogram
GMM(3)
Monolayer
Bilayer
Time(ns)
Z_C.O.M(nm)
Angle (degree)
Rg, Rg, i(i=x,y,z)(nm)
Rg,x
Rg,y
Rg,z
Rg
Dipole – Moment and their components (Debye)
|M|
Mx
My
Mz
Cosine of the angle of Dipole Moment with Z axis, cosϕ

Fig. S21

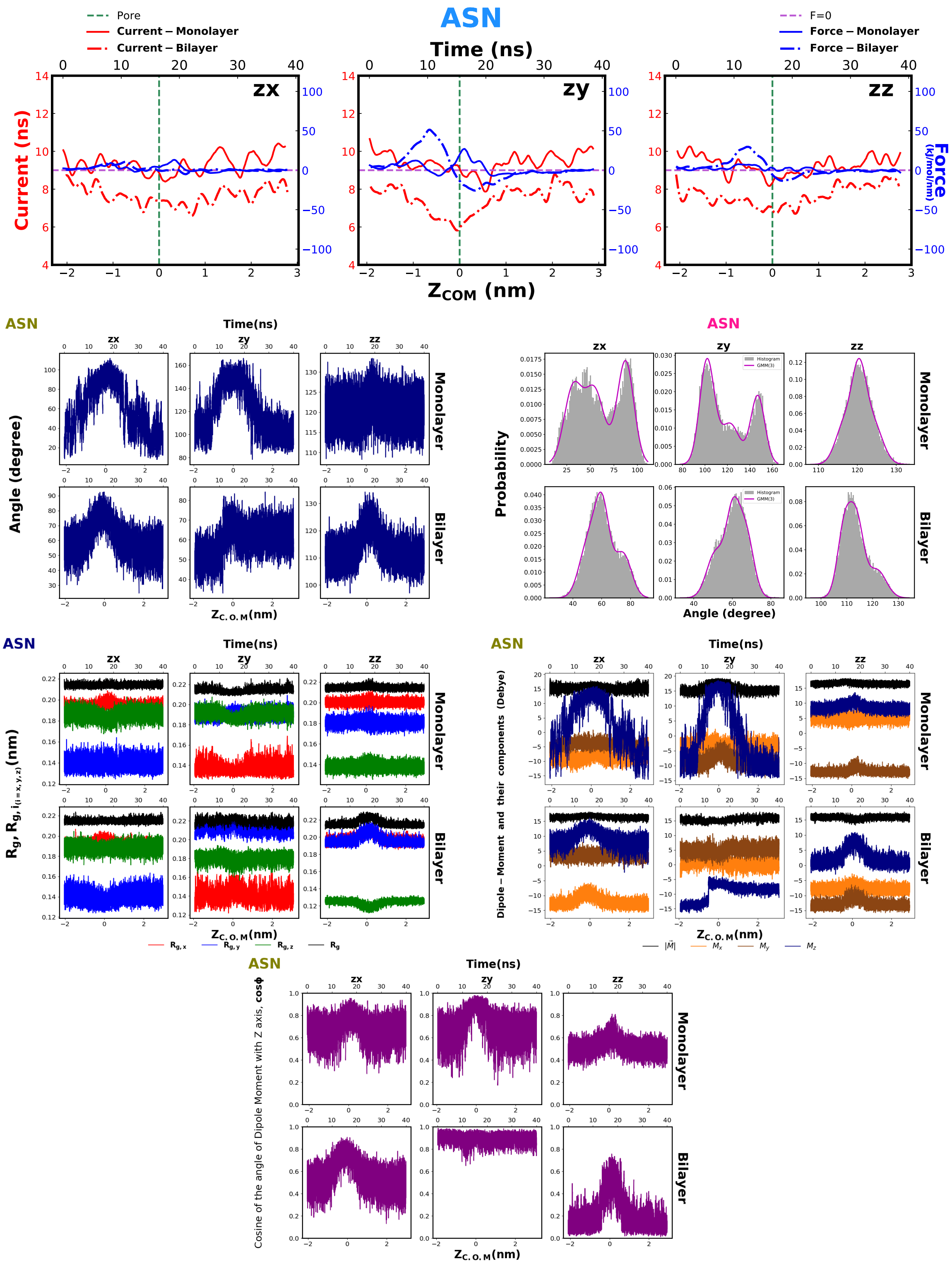
ASN
Pore
Current – Monolayer
Current – Bilayer
F=0
Force – Monolayer
Force – Bilayer
Time (ns)
Current (ns)
Force (kJ/mol/nm)
zx
zy
zz
Z_COM (nm)
Angle (degree)
Monolayer
Bilayer
Z_C.O.M(nm)
Probability
Histogram
GMM(3)
Angle (degree)
R_g, R_g, i(i=x,y,z) (nm)
R_g,x
R_g,y
R_g,z
R_g
Dipole – Moment and their components (Debye)
|M|
M_x
M_y
M_z
Cosine of the angle of Dipole Moment with Z axis, cosϕ

Fig. S22

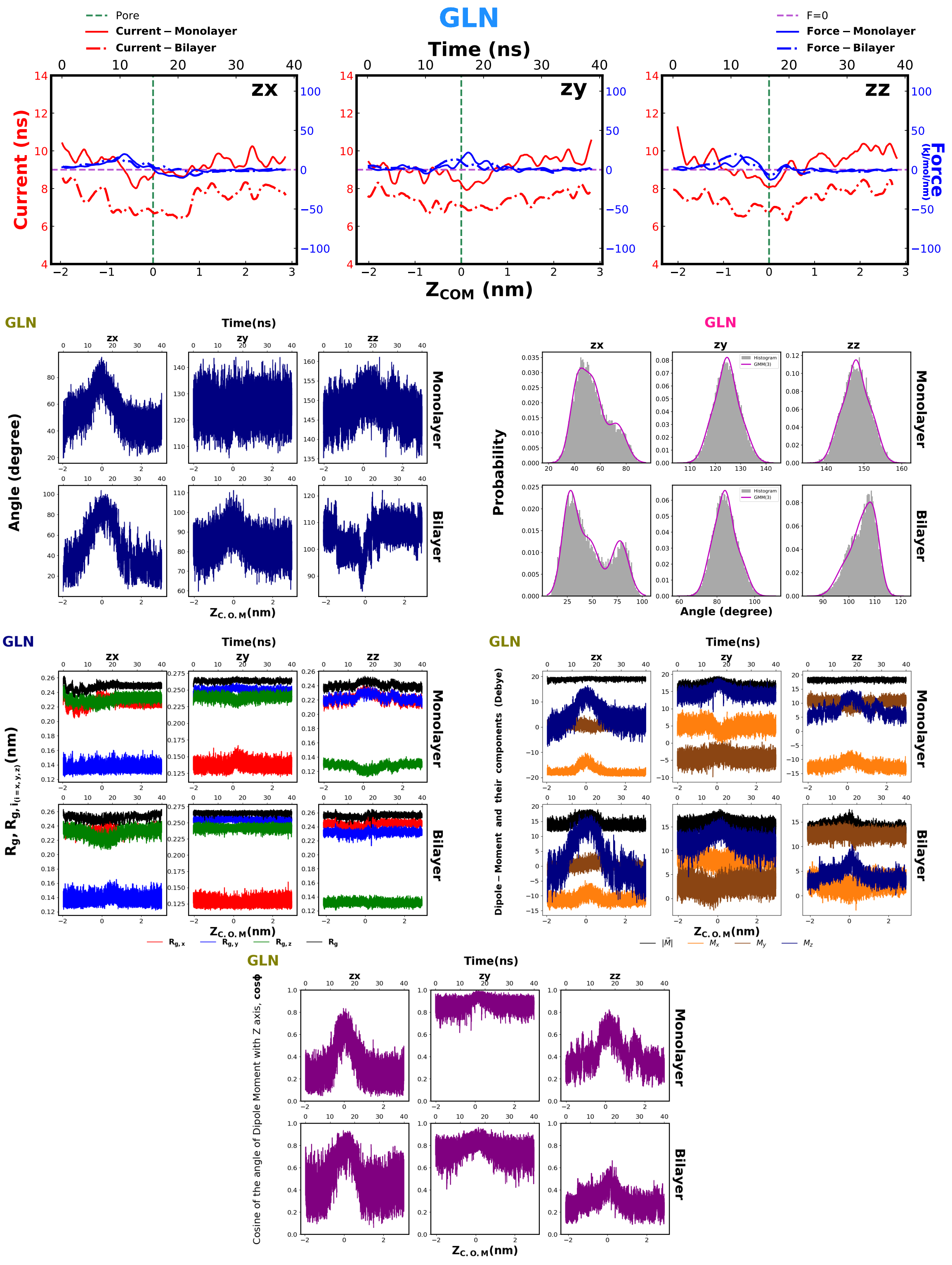

GLN
Pore
Current – Monolayer
Current – Bilayer
F=0
Force – Monolayer
Force – Bilayer
Time (ns)
zx
zy
zz
Current (ns)
Force (kJ/mol/nm)
$Z_{COM}$ (nm)
Angle (degree)
Monolayer
Bilayer
$Z_{C.O.M}$(nm)
Probability
Histogram
GMM(3)
Angle (degree)
$R_g$, $R_{g,i_{(i=x,y,z)}}$(nm)
$R_{g,x}$
$R_{g,y}$
$R_{g,z}$
$R_g$
Dipole – Moment and their components (Debye)
$|\vec{M}|$
$M_x$
$M_y$
$M_z$
Cosine of the angle of Dipole Moment with Z axis, cosϕ

Fig. S23

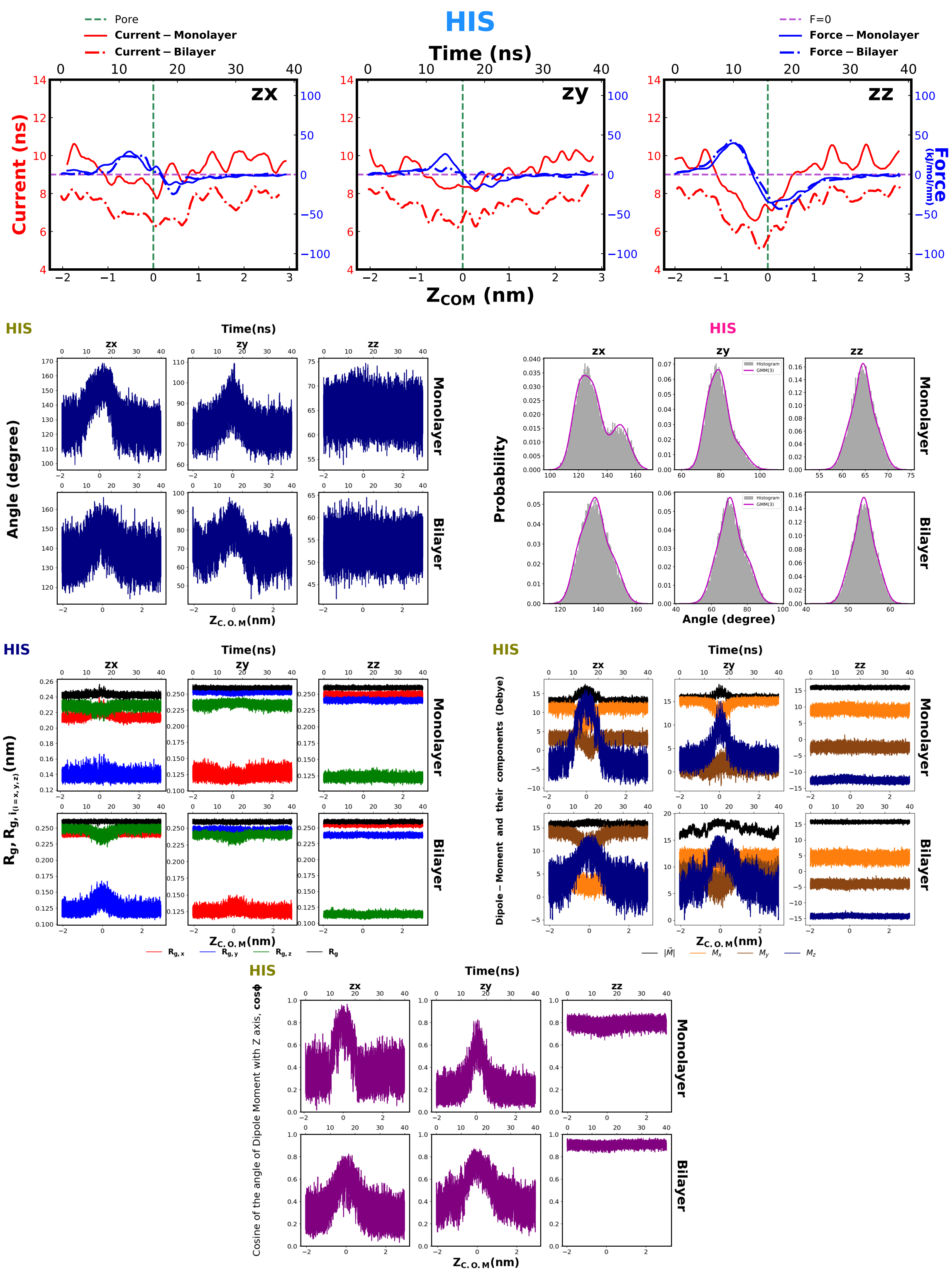
HIS
Pore
Current – Monolayer
Current – Bilayer
F=0
Force – Monolayer
Force – Bilayer
Time (ns)
zx
zy
zz
Current (ns)
Force (kJ/mol/nm)
ZCOM (nm)
HIS
Time(ns)
Angle (degree)
Monolayer
Bilayer
ZC.O.M(nm)
HIS
Probability
Histogram
GMM(3)
Angle (degree)
HIS
Time(ns)
Rg, Rg, i(i=x,y,z)(nm)
Rg,x
Rg,y
Rg,z
Rg
HIS
Time(ns)
Dipole – Moment and their components (Debye)
|M|
Mx
My
Mz
HIS
Time(ns)
Cosine of the angle of Dipole Moment with Z axis, cosϕ
ZC.O.M(nm)

Fig. S24

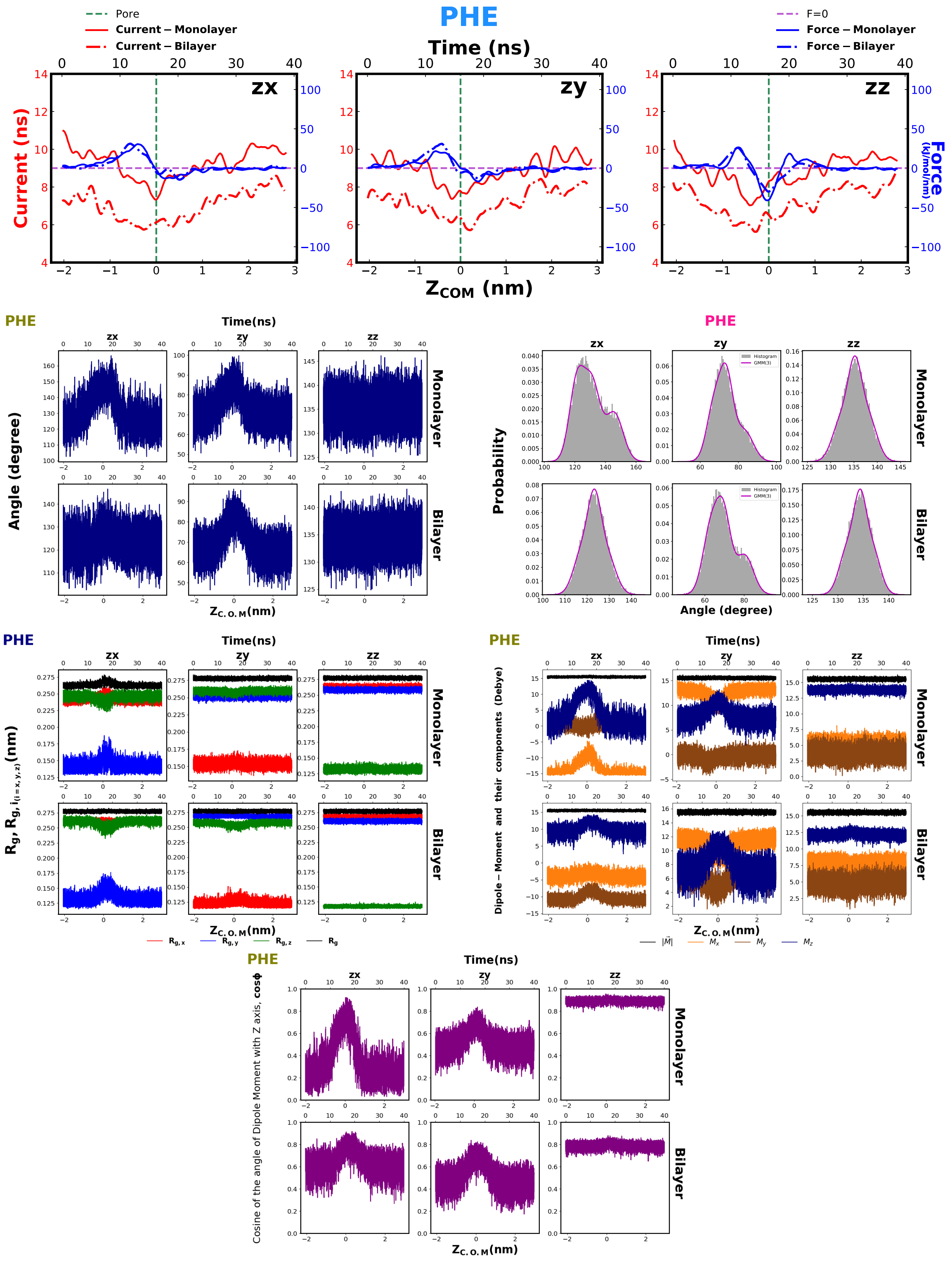

Pore
Current – Monolayer
Current – Bilayer
PHE
Time (ns)
F=0
Force – Monolayer
Force – Bilayer
zx
zy
zz
Current (ns)
Force (kJ/mol/nm)
$Z_{COM}$ (nm)
PHE
Time(ns)
Angle (degree)
Monolayer
Bilayer
$Z_{C.O.M}$(nm)
PHE
Probability
Histogram
GMM(3)
Angle (degree)
PHE
Time(ns)
$R_g$, $R_{g,i(i=x,y,z)}$(nm)
$Z_{C.O.M}$(nm)
$R_{g,x}$
$R_{g,y}$
$R_{g,z}$
$R_g$
PHE
Time(ns)
Dipole – Moment and their components (Debye)
$Z_{C.O.M}$(nm)
$|\vec{M}|$
$M_x$
$M_y$
$M_z$
PHE
Time(ns)
Cosine of the angle of Dipole Moment with Z axis, cosϕ
$Z_{C.O.M}$(nm)

Fig. S25

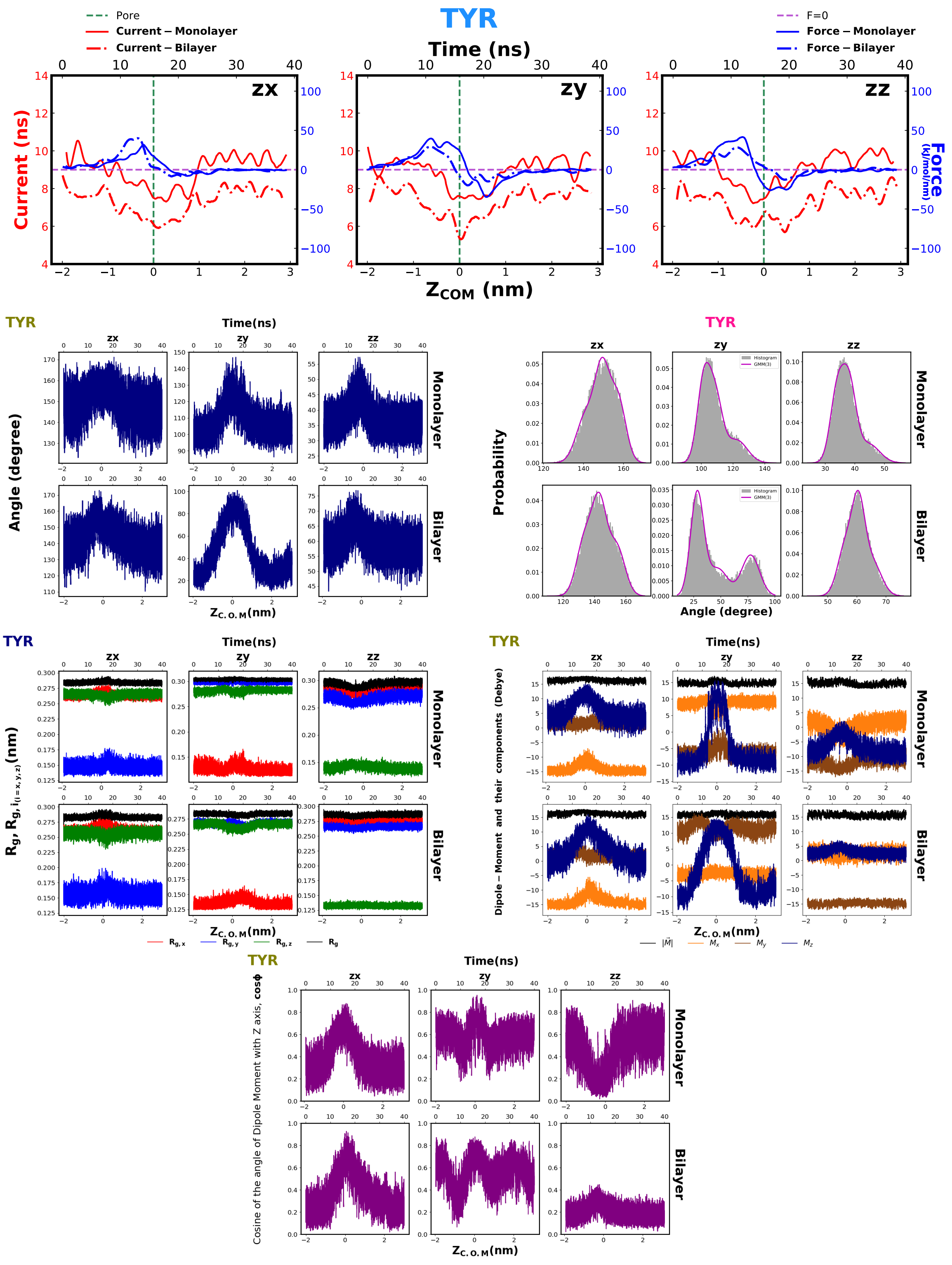
TYR
Pore
Current – Monolayer
Current – Bilayer
F=0
Force – Monolayer
Force – Bilayer
Time (ns)
zx
zy
zz
Current (ns)
Force (kJ/mol/nm)
$Z_{COM}$ (nm)
Time(ns)
Angle (degree)
Monolayer
Bilayer
$Z_{C.O.M}$(nm)
Probability
Histogram
GMM(3)
$R_g, R_{g,i_{(i=x,y,z)}}$(nm)
$R_{g,x}$
$R_{g,y}$
$R_{g,z}$
$R_g$
Dipole – Moment and their components (Debye)
$|\vec{M}|$
$M_x$
$M_y$
$M_z$
Cosine of the angle of Dipole Moment with Z axis, cosϕ

Fig. S26

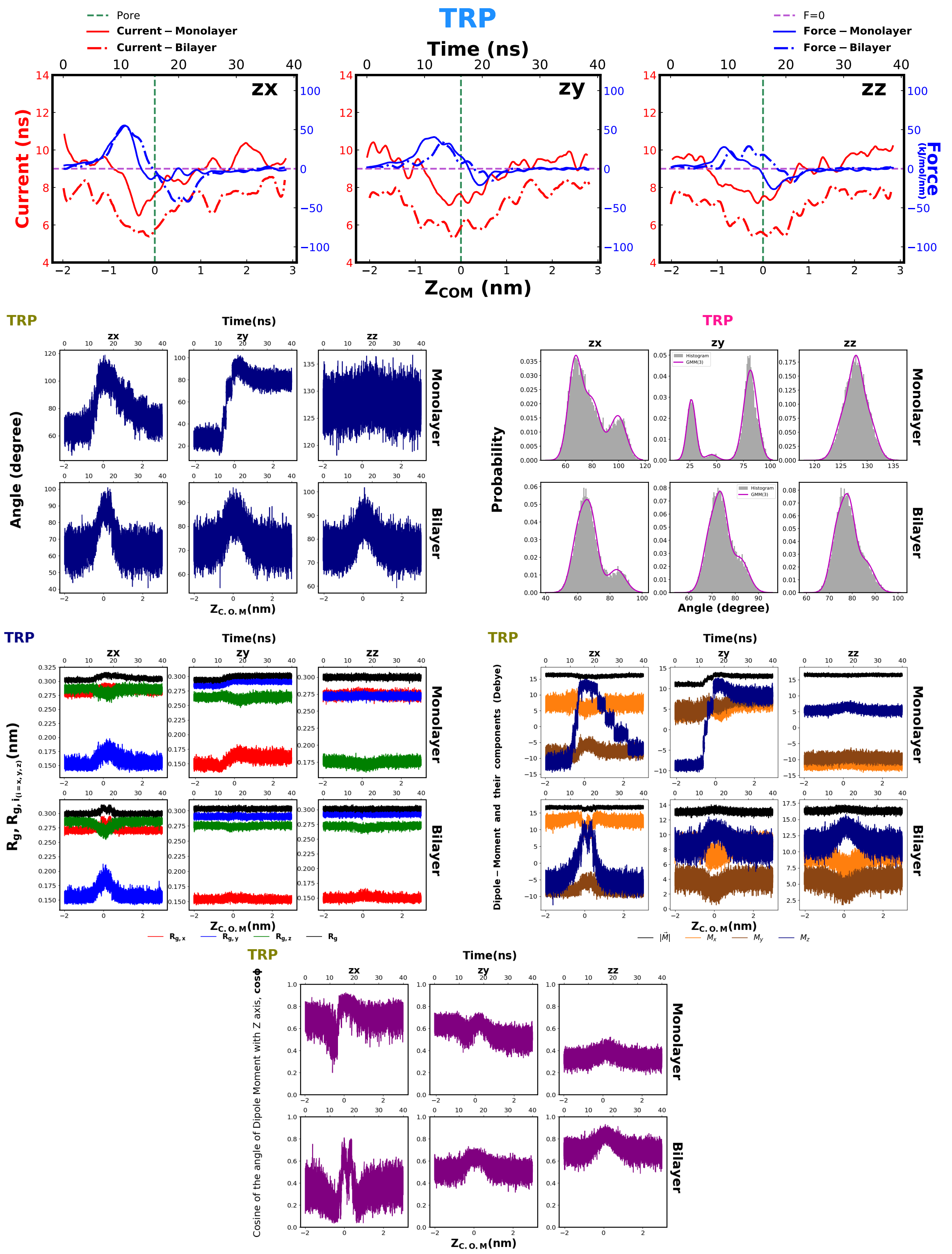

TRP
Pore
Current – Monolayer
Current – Bilayer
F=0
Force – Monolayer
Force – Bilayer
Time (ns)
zx
zy
zz
Current (ns)
Force (kJ/mol/nm)
$Z_{COM}$ (nm)
Angle (degree)
Monolayer
Bilayer
$Z_{C.O.M}$(nm)
Probability
Histogram
GMM(3)
$R_g$, $R_{g,i(i=x,y,z)}$(nm)
$R_{g,x}$
$R_{g,y}$
$R_{g,z}$
$R_g$
Dipole – Moment and their components (Debye)
$|\vec{M}|$
$M_x$
$M_y$
$M_z$
Cosine of the angle of Dipole Moment with Z axis, cosϕ

Fig. S27

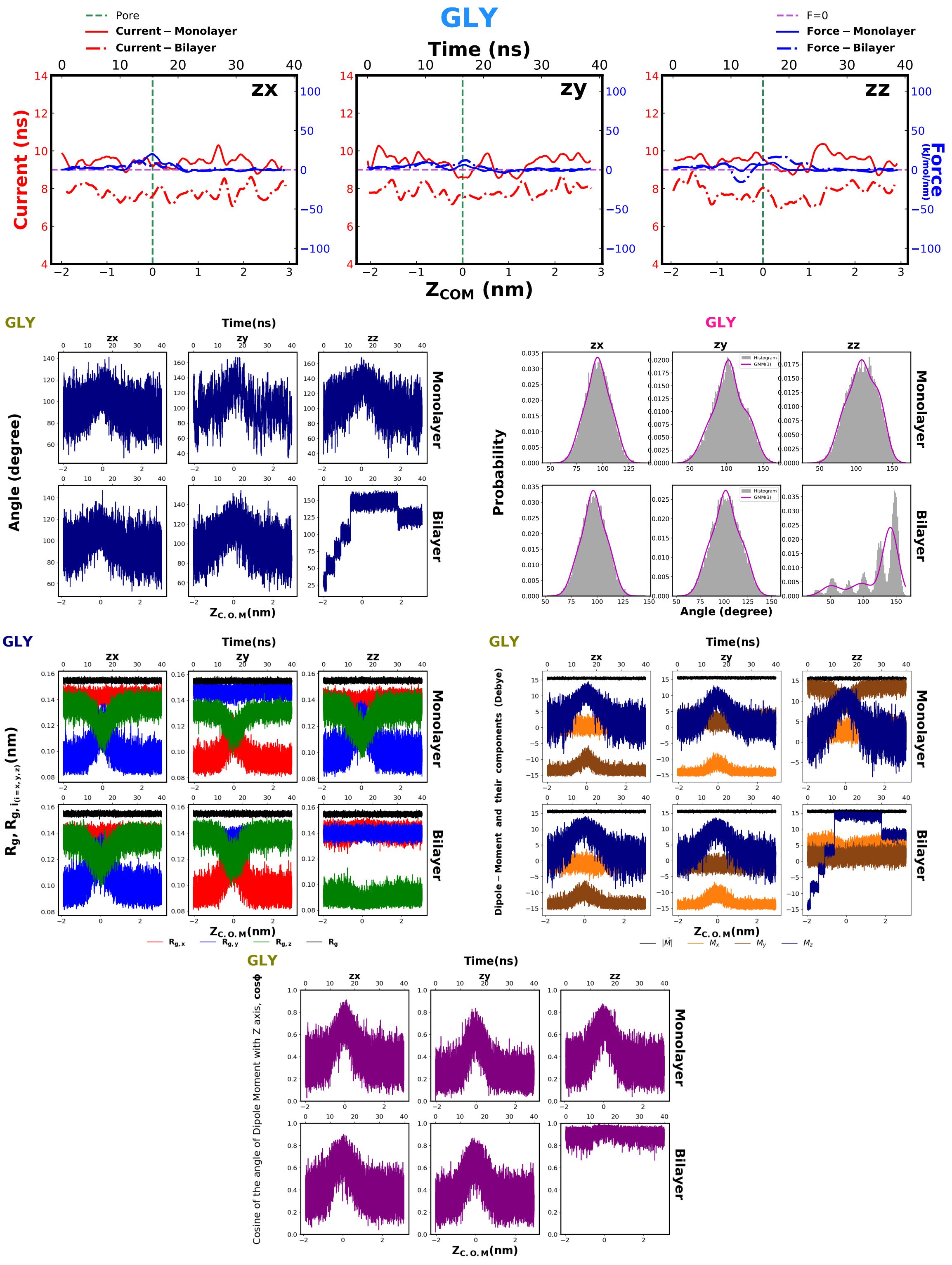
GLY
Pore
Current – Monolayer
Current – Bilayer
F=0
Force – Monolayer
Force – Bilayer
Time (ns)
zx
zy
zz
Current (ns)
Force (kJ/mol/nm)
ZCOM (nm)
Time(ns)
Angle (degree)
Monolayer
Bilayer
ZC.O.M(nm)
Probability
Histogram
GMM(3)
Angle (degree)
Rg, Rg,i(i=x,y,z)(nm)
Rg,x
Rg,y
Rg,z
Rg
Dipole – Moment and their components (Debye)
|M|
Mx
My
Mz
Cosine of the angle of Dipole Moment with Z axis, cosϕ

Fig. S28

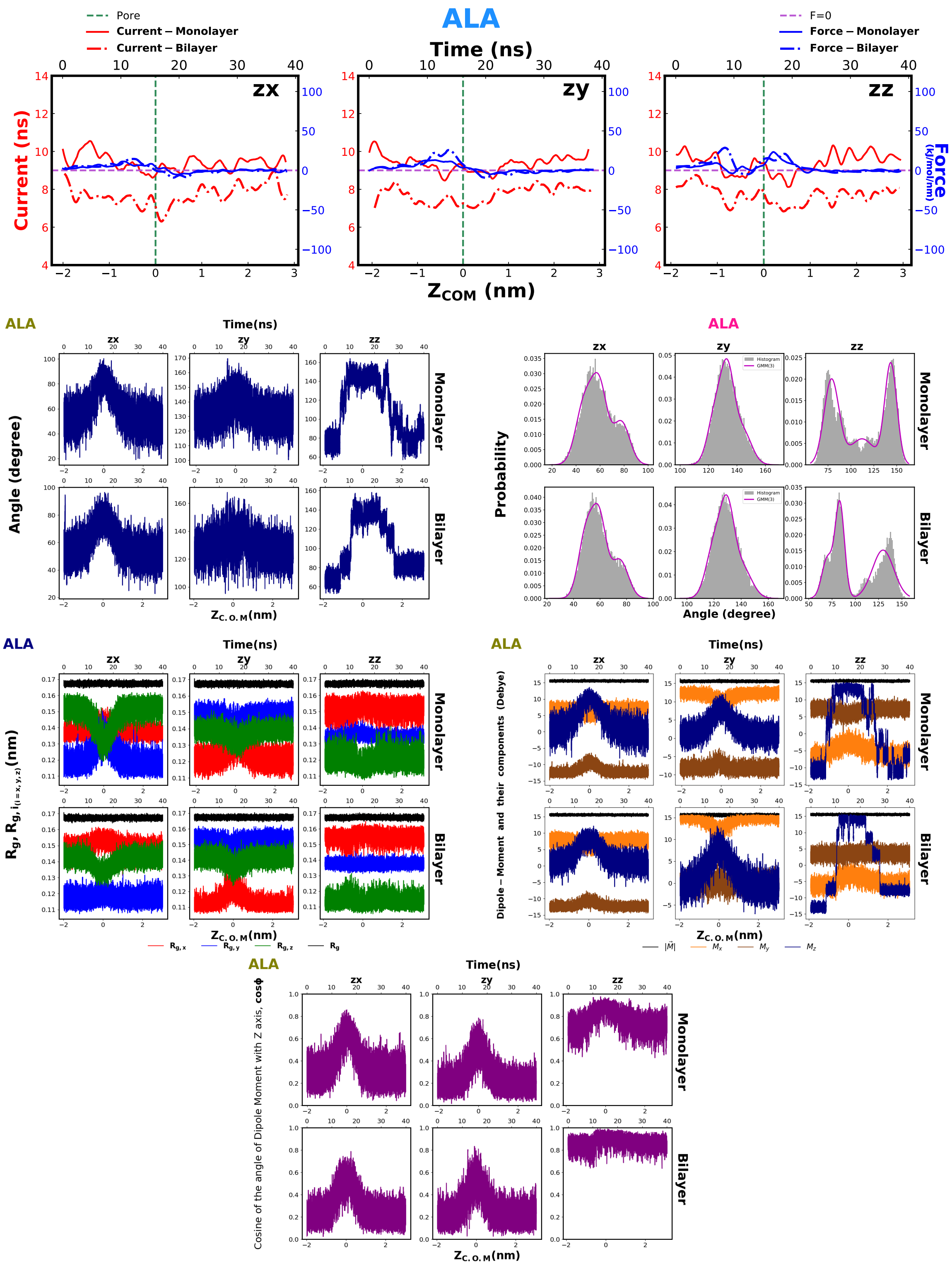
ALA
Pore
Current − Monolayer
Current − Bilayer
F=0
Force − Monolayer
Force − Bilayer
Time (ns)
zx
zy
zz
Current (ns)
Force (kJ/mol/nm)
Z_COM (nm)
Time(ns)
Angle (degree)
Monolayer
Bilayer
Z_C.O.M(nm)
Probability
Histogram
GMM(3)
Angle (degree)
Rg, Rg, i(i=x,y,z)(nm)
Rg,x
Rg,y
Rg,z
Rg
Dipole − Moment and their components (Debye)
|M|
Mx
My
Mz
Cosine of the angle of Dipole Moment with Z axis, cosϕ

Fig. S29

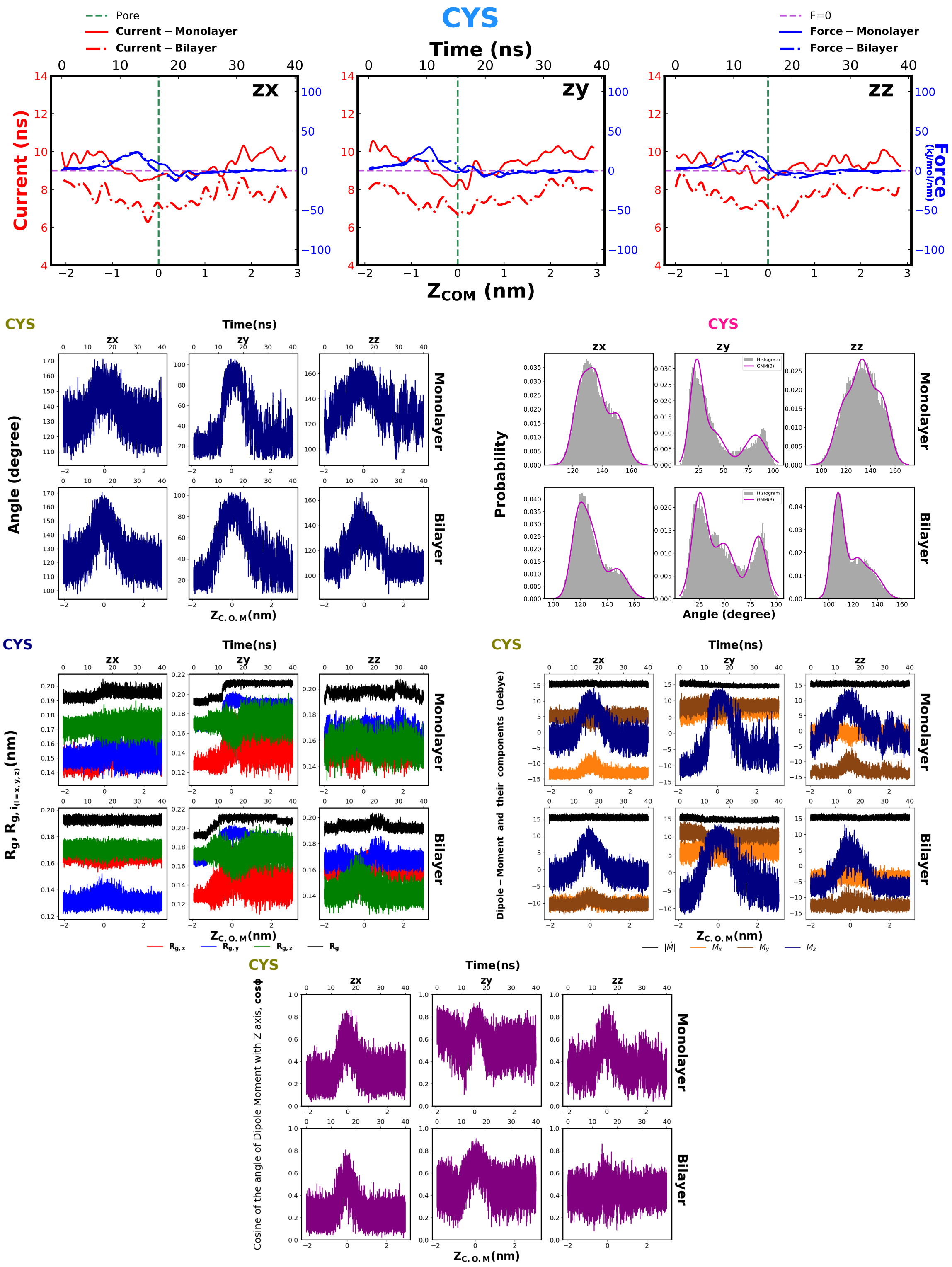
CYS
Pore
Current – Monolayer
Current – Bilayer
F=0
Force – Monolayer
Force – Bilayer
Time (ns)
zx
zy
zz
Current (ns)
Force (kJ/mol/nm)
$Z_{COM}$ (nm)
Angle (degree)
Monolayer
Bilayer
Time(ns)
$Z_{C.O.M}$(nm)
Probability
Histogram
GMM(3)
$R_g$, $R_{g,i_{(i=x,y,z)}}$(nm)
$R_{g,x}$
$R_{g,y}$
$R_{g,z}$
$R_g$
Dipole – Moment and their components (Debye)
$|\vec{M}|$
$M_x$
$M_y$
$M_z$
Cosine of the angle of Dipole Moment with Z axis, cosϕ

Fig. S30

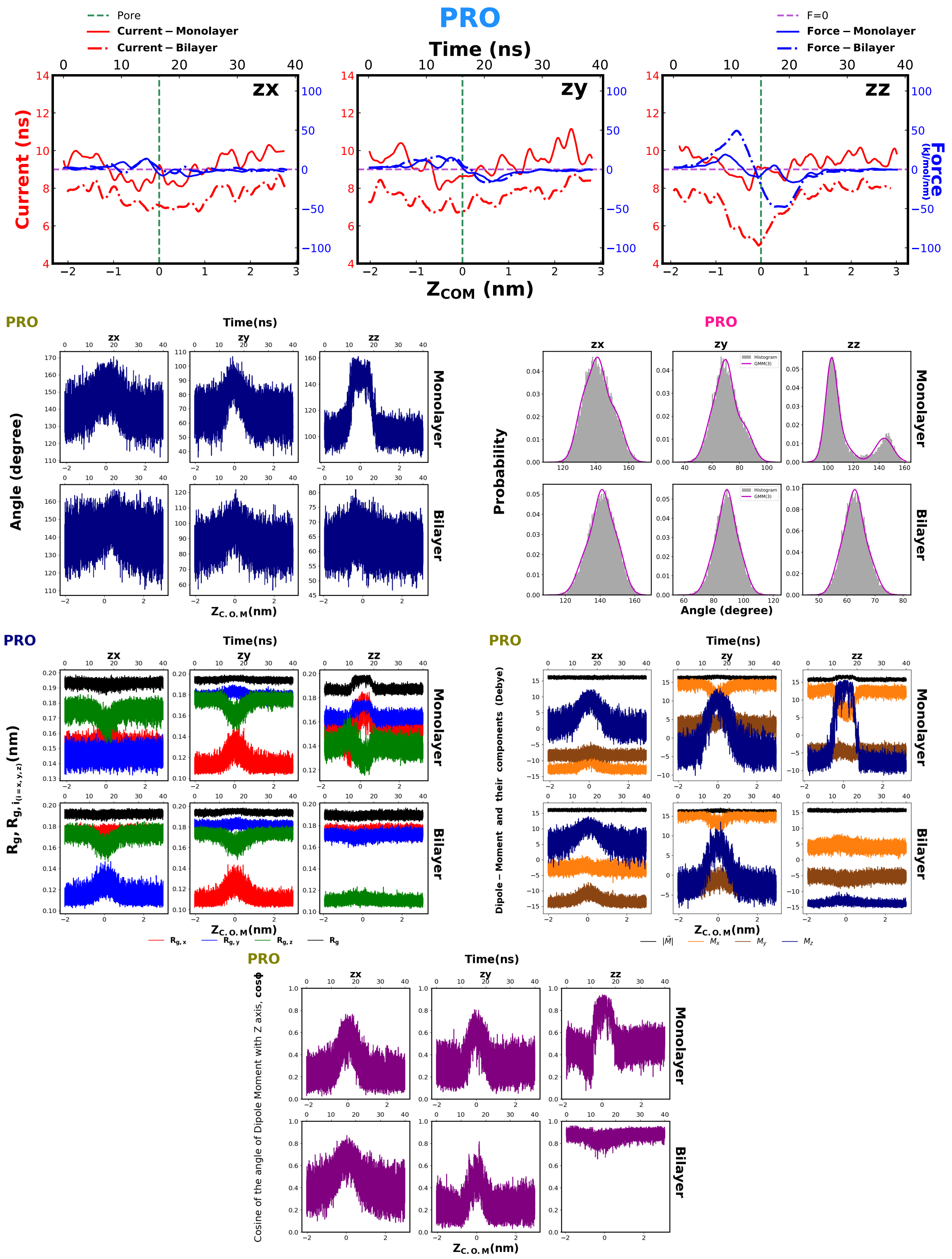

PRO
Pore
Current – Monolayer
Current – Bilayer
F=0
Force – Monolayer
Force – Bilayer
Time (ns)
zx
zy
zz
Current (ns)
Force (kJ/mol/nm)
Z_COM (nm)
Time(ns)
Angle (degree)
Monolayer
Bilayer
Z_C.O.M(nm)
Probability
Histogram
GMM(3)
Angle (degree)
Rg, Rg, i(i=x,y,z)(nm)
Rg,x
Rg,y
Rg,z
Rg
Dipole – Moment and their components (Debye)
|M|
Mx
My
Mz
Cosine of the angle of Dipole Moment with Z axis, cosϕ

Fig. S31

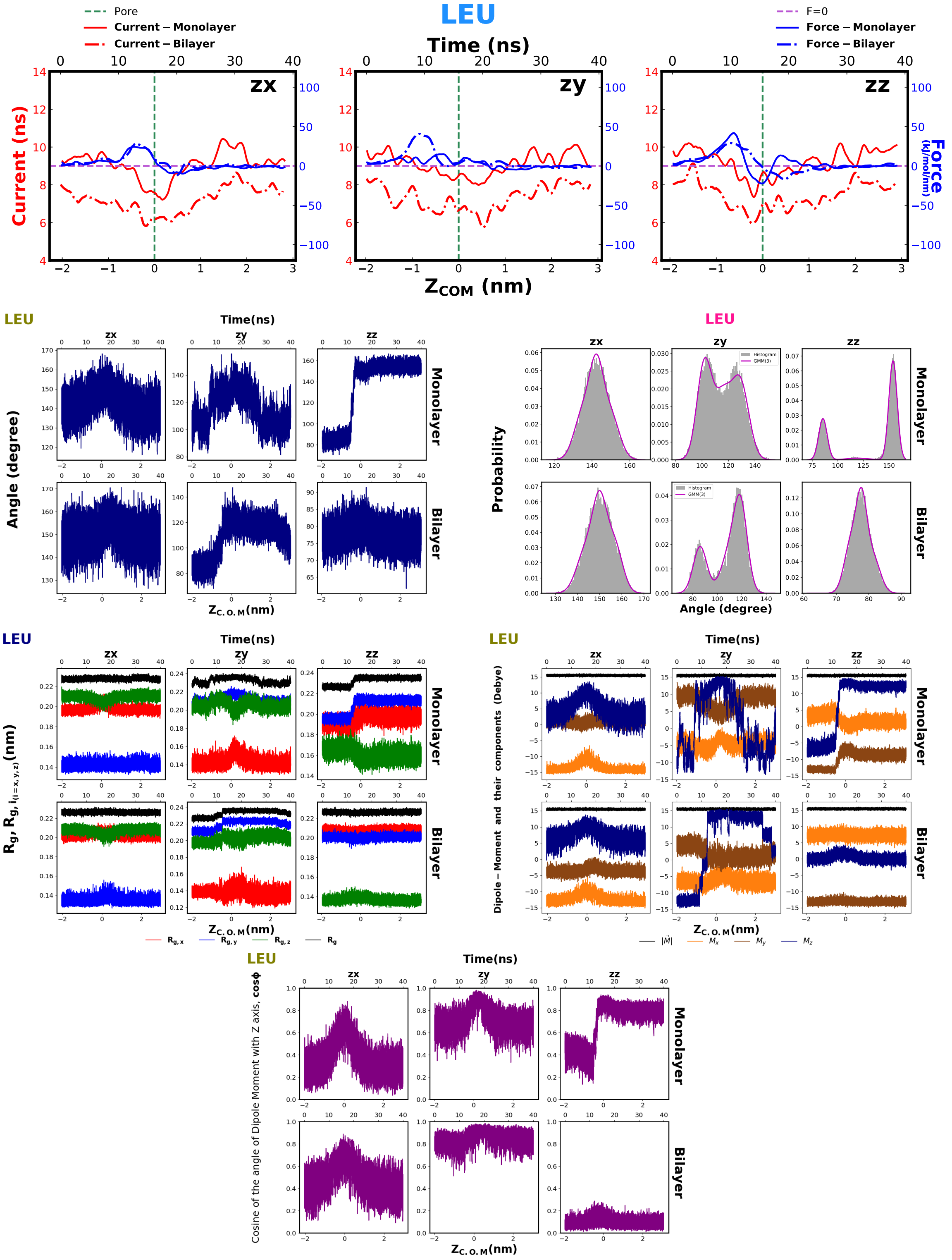
LEU
Pore
Current – Monolayer
Current – Bilayer
F=0
Force – Monolayer
Force – Bilayer
Time (ns)
zx
zy
zz
Current (ns)
Force (kJ/mol/nm)
$Z_{COM}$ (nm)
LEU
Time(ns)
Angle (degree)
Monolayer
Bilayer
$Z_{C.O.M}$(nm)
LEU
Probability
Histogram
GMM(3)
Angle (degree)
LEU
Time(ns)
$R_g$, $R_{g,i_{(i=x,y,z)}}$(nm)
$R_{g,x}$
$R_{g,y}$
$R_{g,z}$
$R_g$
LEU
Time(ns)
Dipole – Moment and their components (Debye)
$|\vec{M}|$
$M_x$
$M_y$
$M_z$
LEU
Time(ns)
Cosine of the angle of Dipole Moment with Z axis, cosϕ

Fig. S32

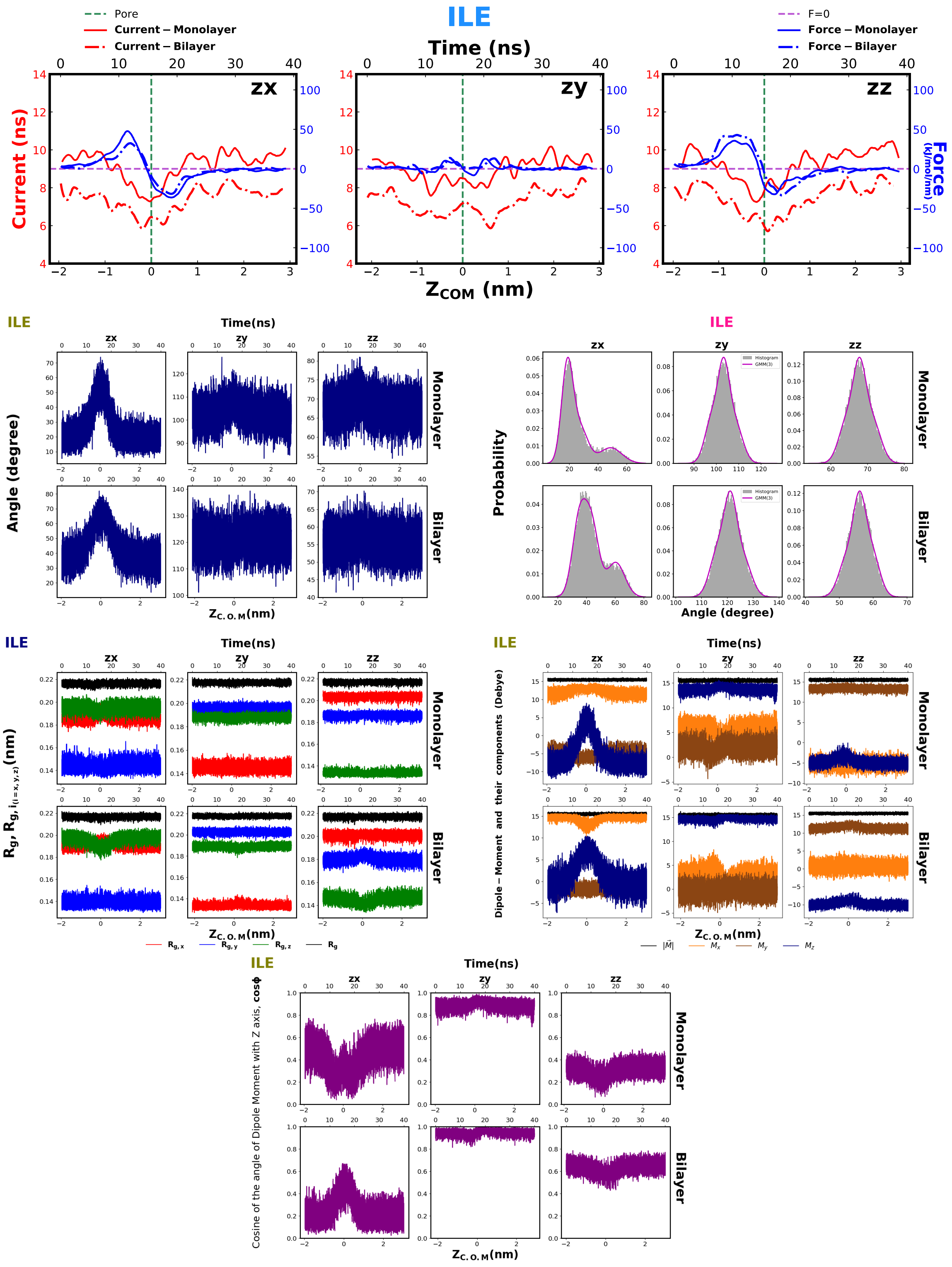
ILE
Pore
Current – Monolayer
Current – Bilayer
F=0
Force – Monolayer
Force – Bilayer
Time (ns)
zx
zy
zz
Current (ns)
Force (kJ/mol/nm)
$Z_{COM}$ (nm)
Angle (degree)
Monolayer
Bilayer
Probability
Histogram
GMM(3)
$Z_{C.O.M}$(nm)
$R_g$, $R_{g,i(i=x,y,z)}$(nm)
$R_{g,x}$
$R_{g,y}$
$R_{g,z}$
$R_g$
Dipole – Moment and their components (Debye)
$|\vec{M}|$
$M_x$
$M_y$
$M_z$
Cosine of the angle of Dipole Moment with Z axis, cosϕ

Fig. S33

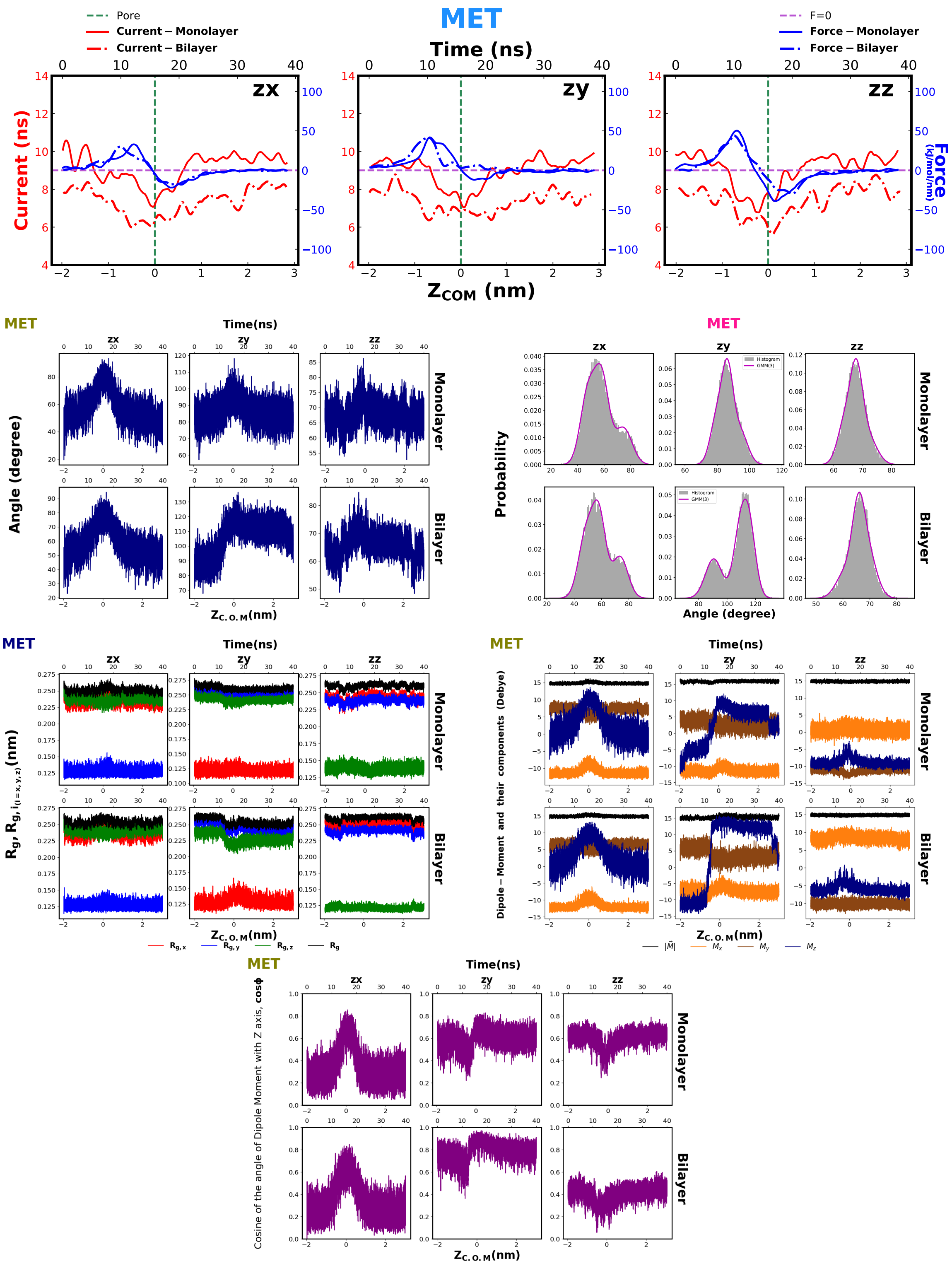
MET
Pore
Current – Monolayer
Current – Bilayer
F=0
Force – Monolayer
Force – Bilayer
Time (ns)
zx
zy
zz
Current (ns)
Force (kJ/mol/nm)
$Z_{COM}$ (nm)
Time(ns)
Angle (degree)
Monolayer
Bilayer
$Z_{C.O.M}$(nm)
Probability
Histogram
GMM(3)
$R_g$, $R_{g,i_{(i=x,y,z)}}$(nm)
$R_{g,x}$
$R_{g,y}$
$R_{g,z}$
$R_g$
Dipole – Moment and their components (Debye)
$|\vec{M}|$
$M_x$
$M_y$
$M_z$
Cosine of the angle of Dipole Moment with Z axis, cosφ

Fig. S34

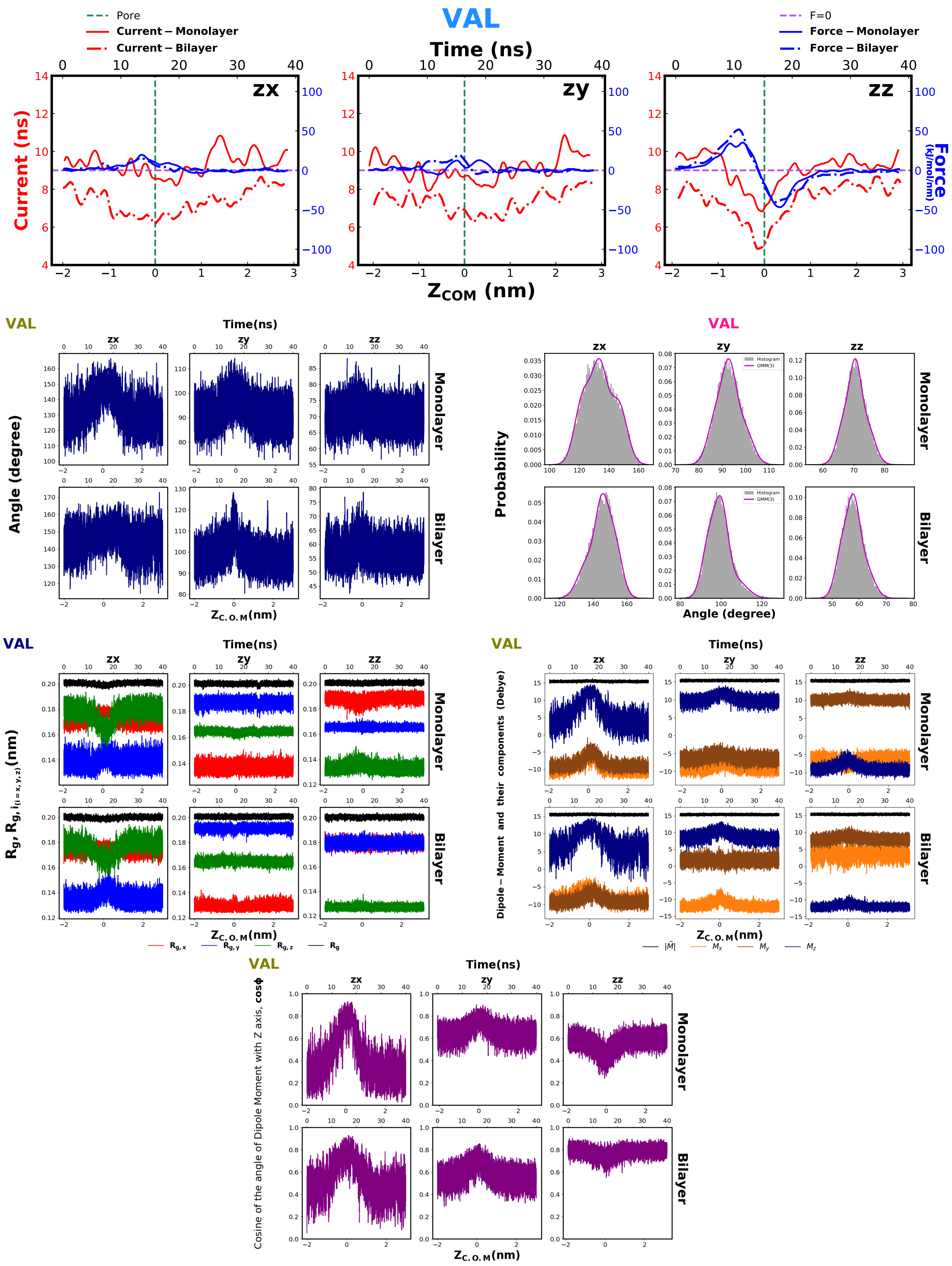
VAL
Pore
Current – Monolayer
Current – Bilayer
F=0
Force – Monolayer
Force – Bilayer
Time (ns)
zx
zy
zz
Current (ns)
Force (kJ/mol/nm)
$Z_{COM}$ (nm)
Angle (degree)
Monolayer
Bilayer
Probability
Histogram
GMM(3)
$Z_{C.O.M}$(nm)
$R_g, R_{g,i_{(i=x,y,z)}}$(nm)
$R_{g,x}$
$R_{g,y}$
$R_{g,z}$
$R_g$
Dipole – Moment and their components (Debye)
$|\vec{M}|$
$M_x$
$M_y$
$M_z$
Cosine of the angle of Dipole Moment with Z axis, cosϕ

### 10 Stretching of the amino acid during translocation

As per the definition used in the article, we have,

$$R_{g,x} = \left( \frac{\sum_i \left( r_{i,y}^2 + r_{i,z}^2 \right) m_i}{\sum_i m_i} \right)^{\frac{1}{2}} \tag{1}$$

$$R_{g,y} = \left( \frac{\sum_i \left( r_{i,x}^2 + r_{i,z}^2 \right) m_i}{\sum_i m_i} \right)^{\frac{1}{2}} \tag{2}$$

$$R_{g,z} = \left( \frac{\sum_i \left( r_{i,x}^2 + r_{i,y}^2 \right) m_i}{\sum_i m_i} \right)^{\frac{1}{2}} \tag{3}$$

$$R_g = \left( \frac{\sum_i \left( r_{i,x}^2 + r_{i,y}^2 + r_{i,z}^2 \right) m_i}{\sum_i m_i} \right)^{\frac{1}{2}} \tag{4}$$

From equations (1),(2),(3),(4) we get

$$2R_g^2 = R_{g,x}^2 + R_{g,y}^2 + R_{g,z}^2 \tag{5}$$

We measure,

$$\delta = \sqrt{2R_g^2 - (R_{g,x}^2 + R_{g,y}^2 + R_{g,z}^2)} \tag{6}$$

as the measure of deviation described in following plot(Fig.S35) for Proline translocation. The more $\delta$ deviates from zero, the more anisotropic the structure of the molecule. Here in Fig.S35 we notice that the overall value of $\delta$ is one order of magnitude smaller than the corresponding $R_g$

in Fig. S30 (figure in the left of third row). The upward spiking of delta in the $|Z_{C.O.M}| < 1$ nm region, in monolayer zz (Fig.S35 c ) and positive value of $\delta^2$ are indicative of the fact that the molecule is getting elongated during translocation. This can be connected with what we see in Fig.5b of the article since an elongated molecule will have increased radius of gyration and that is what we found during translocation.

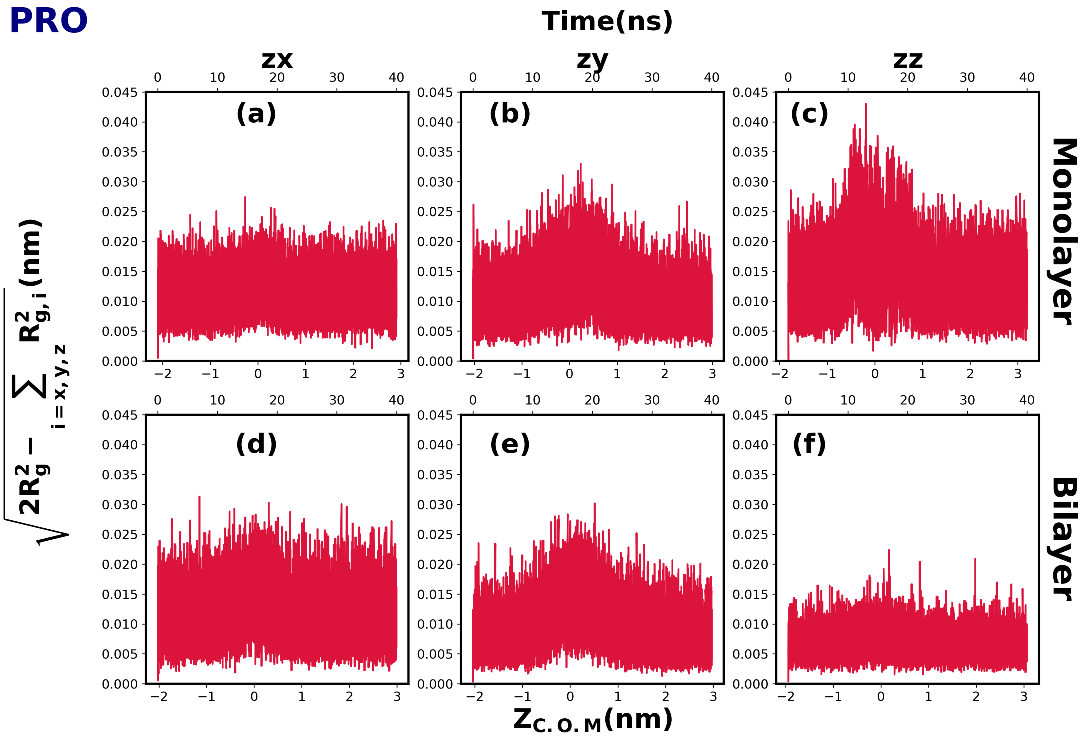


Fig. S35: Stretching of Proline during translocation . The top row (a,b,c) represents monolayer and the bottom one (d,e,f) bilayer. The columns from left to right represent zx (a,b),zy(b,d),zz(c,f) orientation respectively.

### 11 Categorical comparison of Current and Force profile

In this section, we aim to highlight the distinctions in signals combining the current and force, among the members in each subgroups of the amino acids classified as "Charged, Hydrophilic","Neutral, Hydrophobic", "Aromatic, Neutral, Hydrophobic","Neutral, Hydrophobic." We show the results for zz orientation of the amino acids.

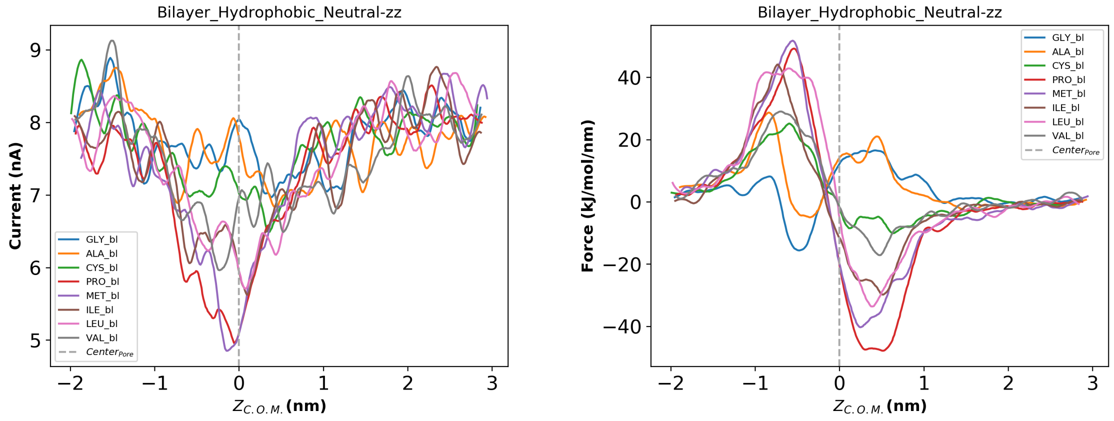


Fig. S36: Neutral, Hydrophobic, Non-aromatic Amino acids. The left panel is for the current signals and the right one is for the force signals. This data is for bilayer. We have shown the same for monolayer in the main article.

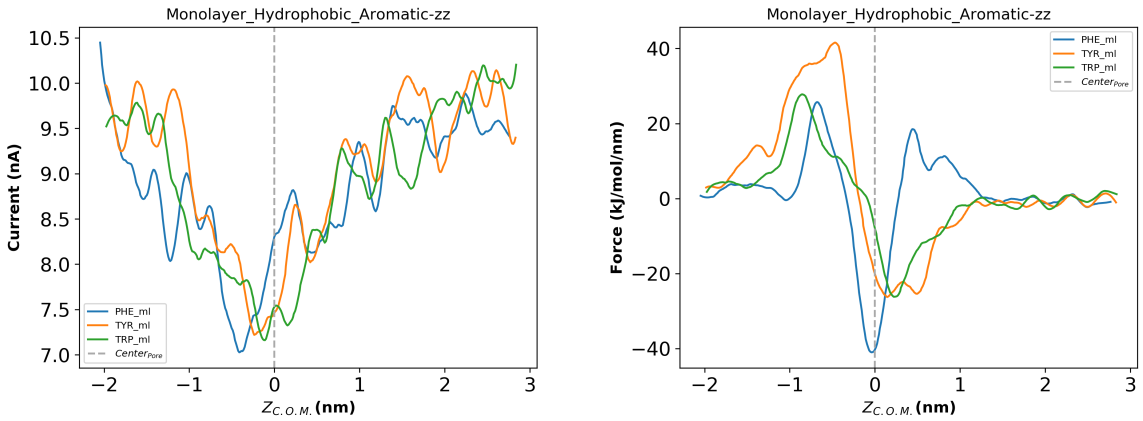


Fig. S37: Aromatic, Neutral, Hydrophobic Amino acids. The left pannel is for the current signals and the right one is for the force signals.The data is for monolayer

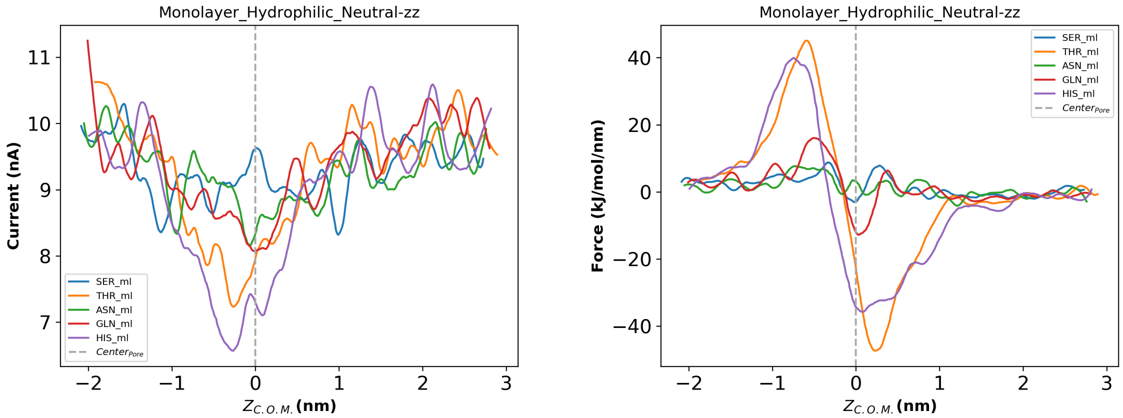


Fig. S38: Neutral,Hydrophilic Amino acids. The left pannel is for the current signals and the right one is for the force signals. The data is for monolayer

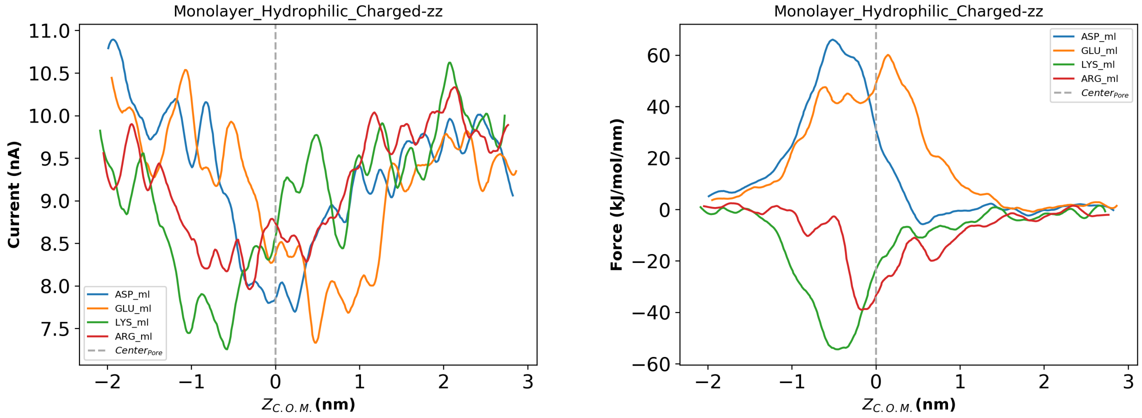


Fig. S39: Charged,Hydrophilic Amino acids. The left pannel is for the current signals and the right one is for the force signals. The data is for monolayer

#### 12 RMSD of $PRO_{zz}$ during translocation.

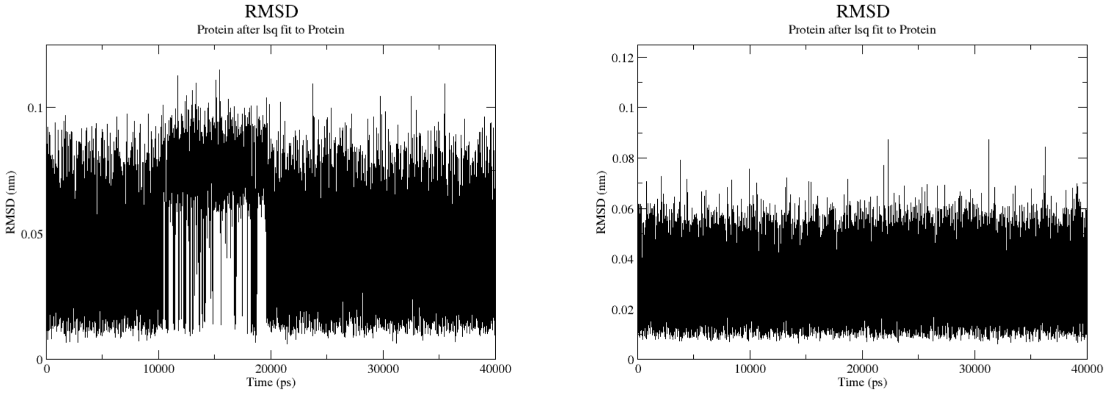


Fig. S40: RMSD profile for Proline translocation in zz orientation for monolayer(left) and bilayer(right)

In the figure above, we calculate the RMSD of the translocating Proline in zz orientation, where the RMSD (root mean square deviation) is calculated from the least squares fit of the structure of translocating amino acid to the initial equilibrated structure of the amino acid. In the left panel, i.e., the monolayer, we see increased structural fluctuation during translocation which corresponds to the flipping we discussed in our manuscript (Fig.5 (b,d,e), Fig.(6)). The right panel shows smaller structural fluctuation overall with no specific increment during translocation, which is the case for bilayer.

### 13 RMSD of the amino acid, comparison of fixed with pulled: Example with $ARG_{zx}$

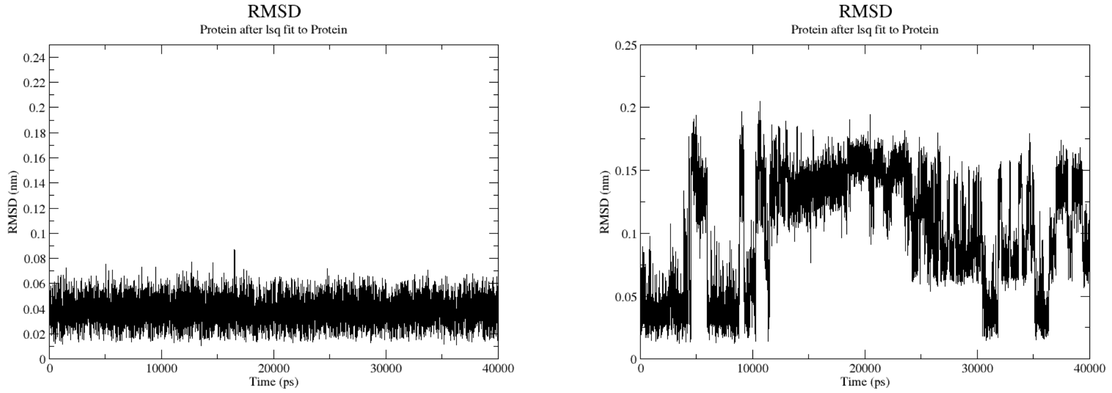


Fig. S41: RMSD of $ARG_{zx}$ while fixed at center compared with RMSD of the same while translocating, for monolayer pore

We can see that for translocating ARG, the RMSD is quite high compared to the static case. This may affect the value of the relative current blockade and possibly be a contributing factor behind the difference in magnitude of relative current blockades we see between static and pulled amino acids.

#### 14 Influence of pore-passivation and adding charge to the membrane on orientation effect

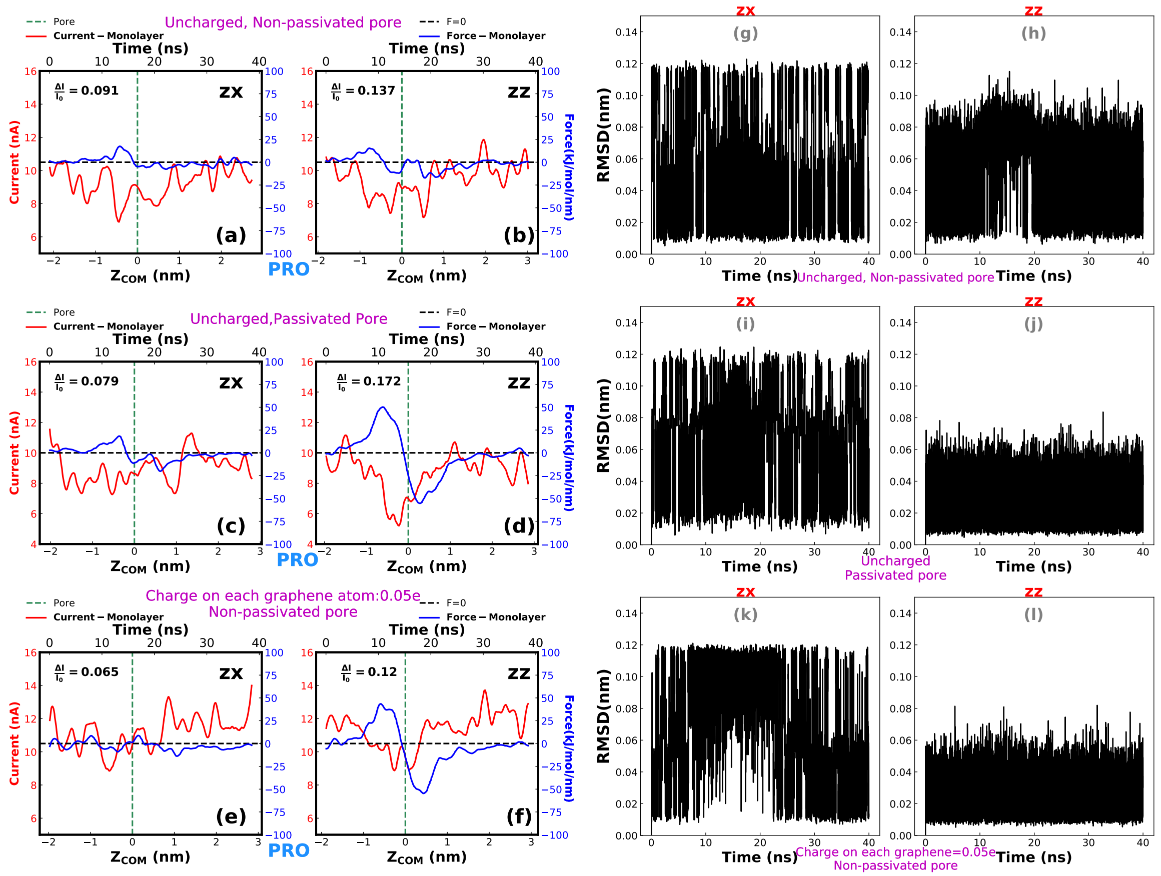


Fig. S42: Current,force and rmsd profiles for PRO translocation in zx and zz orientation. Panel a,b,g,h represents the data for the neutral membrane and pore which was not passivated. This kind of pore was used for the results shown in the article. Panel c,d,i,j represents the data for neutral membrane and pore where the dangling bonds of the pore were passivated with hydrogen atoms. Panel e,f,k,l show the data for the case where every atom of the graphene membrane, including the pore atoms, was charged with +0.05e. The pore remains non-passivated here. We observe that orientation dependence is present in every case.

To investigate the effect of passivation of the pore, we have added hydrogen to the dangling bonds at the pore-mouth (monolayer), where we have used the HP atomtype of charmm36m to model the added hydrogen[7]. The modified pore now looks like what is depicted in Fig.S44. We

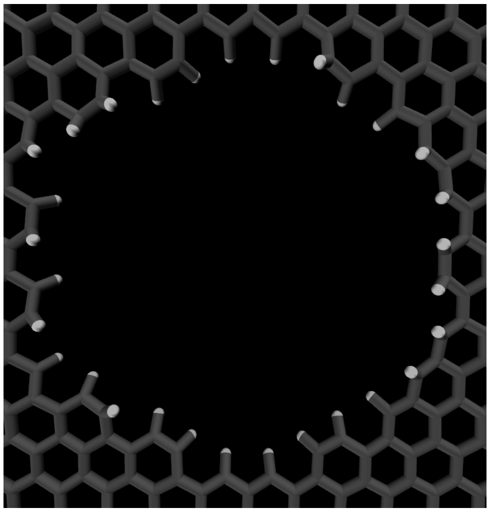

Fig. S43: Hydrogen passivated graphene pore

see in Fig.S43(c,d) that not only the orientation effect exists, it improves over the non-passivated pore(Fig.S43(a,b)) when we compare the relative current blockades of zx and zz orientation.

In the same figure S43, we also show current and force profiles for Proline translocation through a pore in monolayer charged graphene. We have assigned a charge of approximately 0.05e on each carbon atom of the membrane to achieve a charge density of 0.28 $C/m^2$ [8]. Fig.S43 (e,f)demonstrates that the overall current increases for the charged membrane compared to the current shown in Fig.S43(a,b) for PRO translocation through the neutral membrane pore. We can clearly see the difference between the relative current blockade in the zx and zz orientation also in the charged case.

Moreover, in the zz orientation for both the passivated ( Fig.S43(c,d))and the charged pore ( Fig.S43(e,f)), we see that the magnitude of force in zz orientation is higher than zx orientation with a prominent antisymmetric nature about the pore, associated with higher current blockade in zz orientation. For the neutral non-passivated membrane,this kind of behavior was also mostly populated in the zz orientation, shown in Fig.3 (c,f) of the manuscript with square markers. In Fig.S43(g-l), we compare the RMSD of the structure of the amino acid during translocation, for the orientation zx and zz. We find a lower fluctuation in the zz orientation, which in turn produces a better current blockade. This is similar to what we described in the manuscript.

To probe the existence of orientation effect when the amino acid is static at the pore in a charged membrane, we have experimented with GLY, ASN and ARG to cover most of the volume range of the amino acids. Fig.S45 shows that the decrease in current is less sensitive to the increase in volume of the amino acid for orientation zz, which is also what we find for the neutral membrane (Fig. 2 of our manuscript).

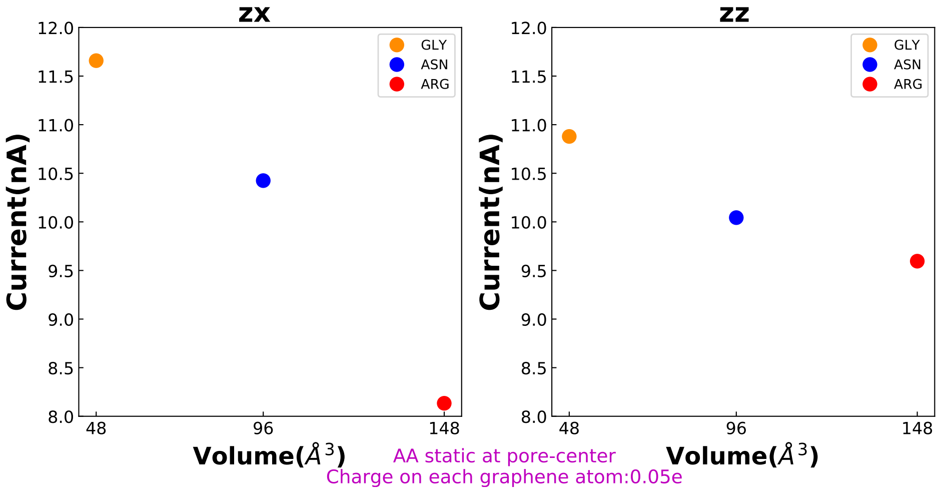


Fig. S44: Volume-Current correlation for amino acids static at charged membrane pore

In conclusion, the effect of orientation of the amino acid in current blockade exists for a charged membrane pore and a passivated pore. However, to obtain a better understanding of the orientation effect subjected to a charged pore and a passivated pore, we need to perform an exhaustive set of simulations involving all the amino acids and their different orientations.

#### 15 Number of atoms and molecules in the simulations.

Table S2

| **Layer No.** | **Water Molecules** | **$K^+$ ions** | **$Cl^-$ ions** | **Graphene atoms** | **Total No. of atoms including the amino acid** |
|---|---|---|---|---|---|
| **Monolayer** | **8286-8293** | **166-167** | **166-167** | **886** | **26091-26119** |
| **Bilayer** | **8182-8190** | **169-170** | **169-170** | **1778** | **26684-26712** |

#### 16 The probability density of currents for different amino acids.

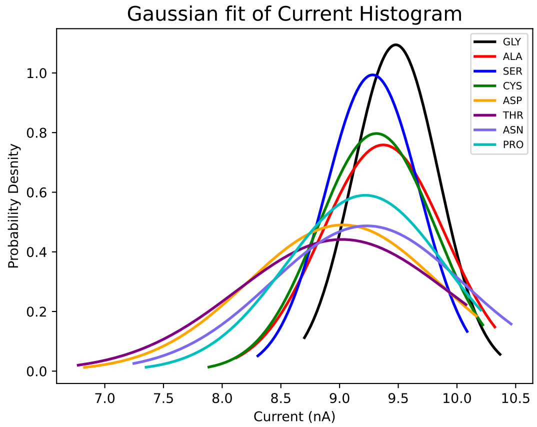


Fig. S45: The probability density of the current plots shown in Fig.4c of the article

#### 17 Structures and Properties of the Twenty Amino Acids

Table S3

| Name | Structure[a] | vdW volume[b] ($\AA^3$) | charge (e) |
|---|---|---|---|
| GLY(G) | 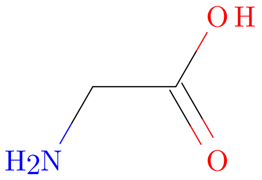 | 48 | 0 |
| ALA(A) | 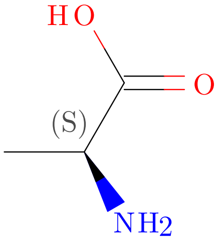 | 67 | 0 |
| CYS(C) | 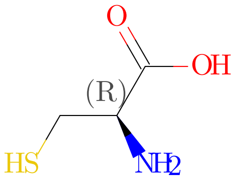 | 86 | 0 |
| PRO(P) | 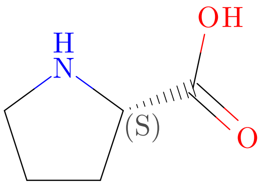 | 90 | 0 |

| **Name** | **Structure**[a] | **vdW volume**[b] (Å$^3$) | **charge (e)** |
|---|---|---|---|
| VAL(V)) | 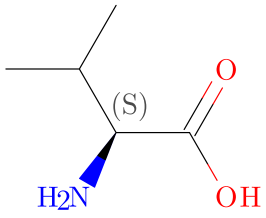 | 105 | 0 |
| MET(M) | 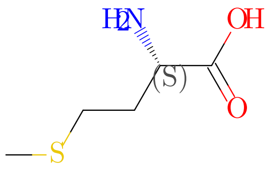 | 124 | 0 |
| LEU(L) | 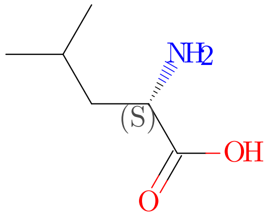 | 124 | 0 |
| ILE(I) | 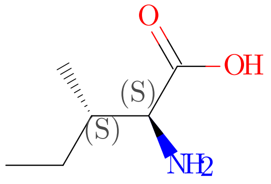 | 124 | 0 |

| **Name** | **Structure**[a] | **vdW volume**[b] ($\AA^3$) | **charge (e)** |
|---|---|---|---|
| SER(S) | 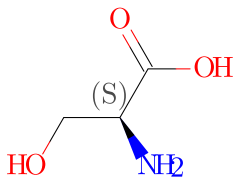 | 73 | 0 |
| THR(T) | 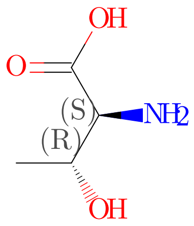 | 93 | 0 |
| ASN(N) | 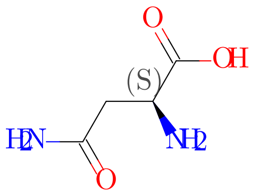 | 96 | 0 |
| GLN(Q) | 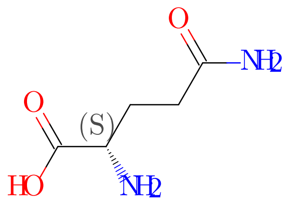 | 114 | 0 |
| HIS(H) | 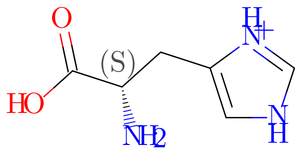 | 118 | 0 |

| Name | Structure[a] | vdW volume[b] ($\AA^3$) | charge (e) |
|---|---|---|---|
| ASP(D) | 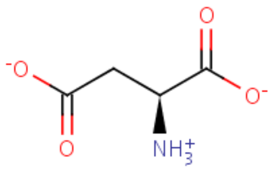 | 91 | -1 |
| GLU(E) | 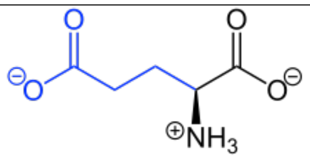 | 109 | -1 |
| LYS(K) | 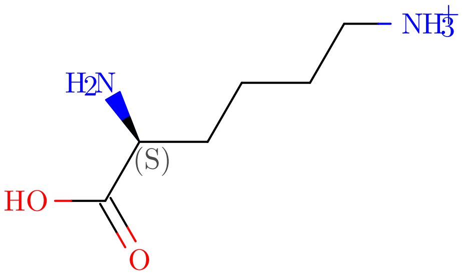 | 135 | +1 |
| ARG(R) | 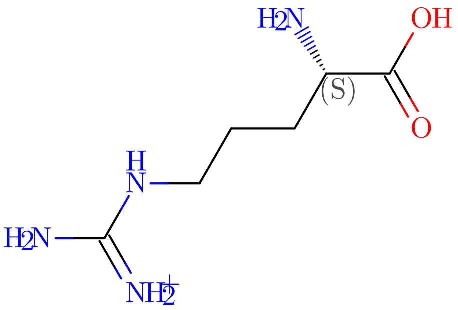 | 148 | +1 |

| **Name** | **Structure**[a] | **vdW volume**[b] ($\AA^3$) | **charge (e)** |
|---|---|---|---|
| PHE(F) | 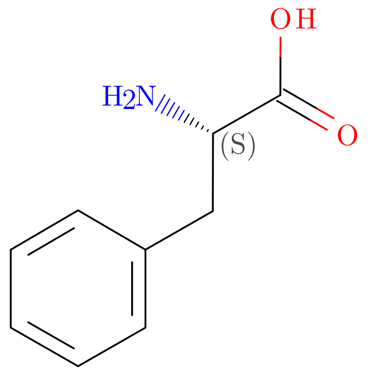 | 135 | 0 |
| TYR(Y) | 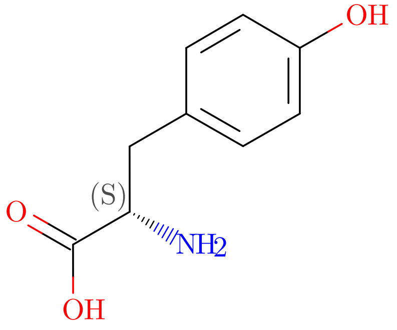 | 141 | 0 |
| TRP(W) | 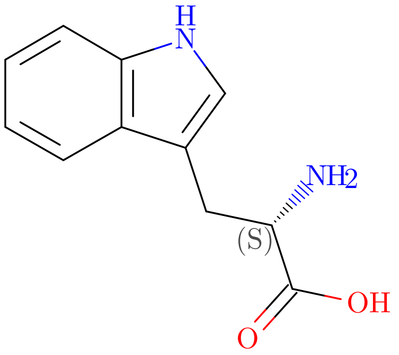 | 163 | 0 |

[a] Structures have been taken from RCSB PDB. Aspertate structure was taken from ChEMBL database and glutamate structure was taken from Wikipedia. [b] Data adapted from Darby and Creighton (1993) [5]
